# Plant Organellar Homologous Recombination Depends on Accessory Factors for Recombinase Activity

**DOI:** 10.64898/2026.10.24.761589

**Authors:** Josué Mora-Garduño, Antolín Peralta-Castro, Andrés Aguilar-Granados, Claudia D. Raygoza, Corina Díaz-Quezada, Luis G. Brieba

**Affiliations:** Cinvestav-Unidad de Genómica Avanzada, Km. 9.6 Libramiento Norte Carretera. Irapuato-León, Irapuato, Guanajuato, México, 36821

## Abstract

Plant mitochondrial and plastid genomes rely on homologous recombination (HR) for error-free DNA repair of double strand breaks. Here, we reconstructed the homologous recombination pathway of plant organelles from Arabidopsis, aiming to understand how proteins of diverse evolutionary origins assemble into a functional repair pathway. We found that the three bacterial-derived recombinases, AtRecAs assemble D-loop intermediates, but only chloroplast AtRecA1 independently execute end-joining recombination on large DNA substrates. AtRecA1-mediated end-joining recombination is enhanced by plant-unique single-stranded DNA-binding proteins of the Whirly and Organellar Single-Stranded Binding (OSB) families, in contrast these proteins are mandatory for the recombination activities of the dual-targeted AtRecA2 and mitochondrial AtRecA3. Recombinational intermediates generated by AtRecAs are resolved by a bacterial-like RadA helicase but not bacterial-like RecG. Finally, AtRecX inhibits the recombinase activities of AtRecA2, while the activities of AtRecA1 and AtRecA3 remains largely insensitive. Collectively, our results provide an integrative reconstitution of the plant organellar homologous recombination machinery, revealing that homologous recombination arises from a complex system of bacterial, phage-derived, plant-specific, and eukaryotic components. Plant-specific ssDNA-binding proteins emerge as key regulators of this process, promoting recombination efficiency and genome stability, that supports the essential functions of homologous recombination in DNA repair and DNA replication maintenance.

**GRAPHICAL ABSTRACT:** 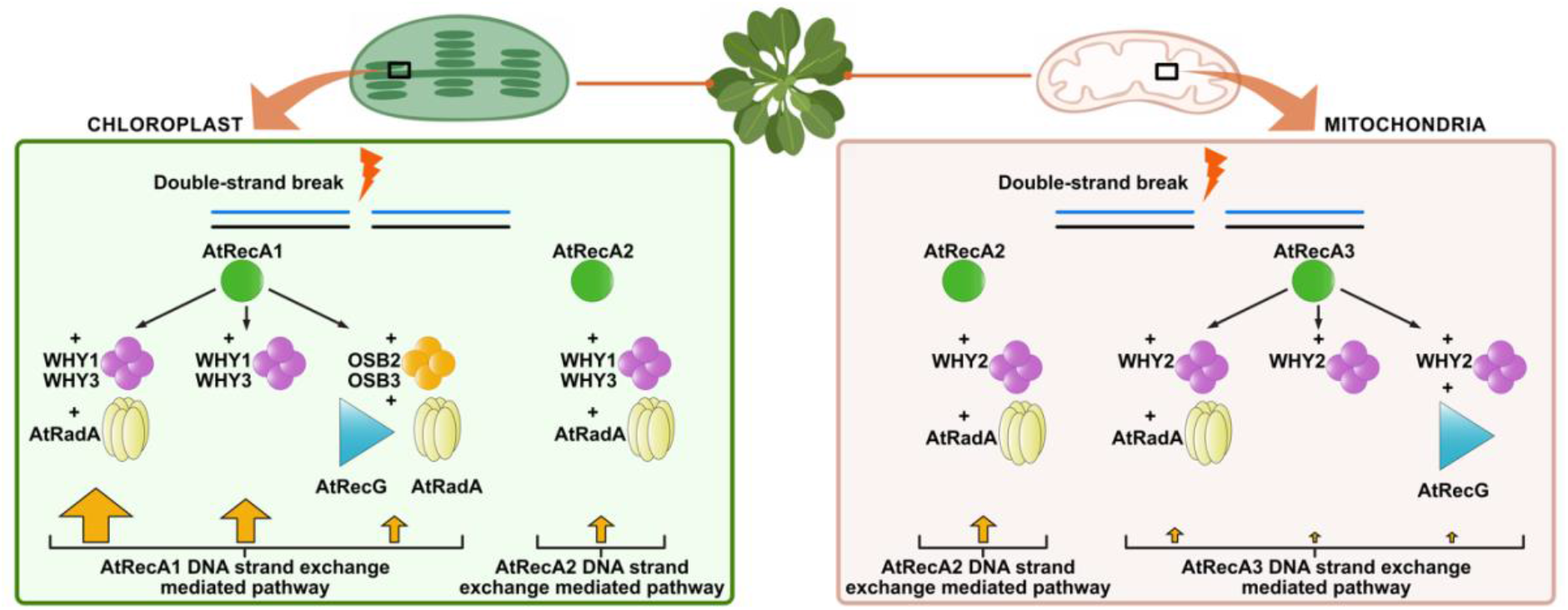

DNA repair pathways mediated by homologous recombination in plant organelles

## INTRODUCTION

Plant organellar genomes are exposed to DNA damaging agents that induce double-stranded DNA breaks (DSB) and genomic rearrangements (1–3). Organisms cope with DBS using at least three different DNA repair routes: Homologous Recombination (HR), non-homologous end joining (NHEJ), and micro-homology mediated end joining (MMEJ). Plant organellar genomes harbor signatures for the enzymatic activity of these three routes (4–10). However, the most characteristic feature of plant organellar genomes is their propensity to recombine (11–13). Unlike the almost absent homologous recombination events in animal mitochondria, plant mitochondrial and chloroplast genomes are highly recombinogenic (14–16). In flowering plants chloroplasts, a “flip-flop” recombination event between large-inverted repeats generates two equimolar plastome isoforms that differ in the orientation of the single-copy regions (17–19). Plant mitochondrial genomes harbor a variety of repeated sequences, from intermediate-site and large repeated sequences (> 1 kb) that frequently recombine. These repeated recombination events give rise to a multipartite, highly redundant genome architecture in which alternative subgenomic molecules coexist at approximately equimolar stoichiometries (20). In addition, homologous recombination events occur sporadically at intermediate-site repeats between 100 to 1000 bp generating rearranged DNA molecules known as sublimons (substoichiometric genomes) (12,21,22). Notably plant organellar genomes harbor an enzymatic machinery able to execute homologous recombination, as seminal studies have predicted the components of the organellar HR pathway in the plant model *Arabidopsis thaliana* (5,6,8,11,22–26). However, these proteins or enzymes have been investigated individually, and their coordinated activities remain poorly understood. The highly recombinogenic nature of plant organellar genomes, together with the mosaic evolutionary origins of their DNA repair factors and their exceptionally low mutation rates, highlights the need to elucidate the molecular basis of homologous recombination in mitochondria and chloroplasts as an integrated system. The low mutation rates of organellar genomes further suggest that the highly accurate HR DNA repair pathways must operate to maintain genome stability (22,27,28). To fill this gap, we subcloned and purified all enzymatic components of the homologous recombination machinery in plant mitochondria and chloroplast identified to date from the plant model *Arabidopsis thaliana*. These factors include single-stranded DNA-binding proteins (SSB1, SSB2, Whirlies, and OSBs), recombinases (RecA1, RecA2, and RecA3), the recombination inhibitor RecX, and DNA helicases RadA and RecG.

## MATERIALS AND METHODS

### Cloning

The genes coding for SSB1 (At4g11060), SSB2 (At3g18580), WHY1 (AT1G14410), WHY2 (At1g71260), WHY3 (AT2G02740), OSB1 (At1g47720), OSB2 (At4g20010), OSB3 (At5g44785), AtRadA (At5g50340) and AtRecG (At2g01440.1) lacking their predicted targeting signals were amplified from Arabidopsis cDNA and cloned in bacterial expression vectors as indicated in **Table S1**. The genes coding for OSB4 (At1g31010), AtRecA1 (AT1G79050), AtRecA2 (At2g19490), AtRecA3 (At3g10140), and AtRecX (At3g13226.1) were coding optimized for their expression in bacteria (Twist Bioscience, California, USA) and cloned in bacterial expression vectors (**Table S1**).

### Construction of point and deletion mutants

Point mutants were constructed using the_Q5^®^ Site-Directed Mutagenesis Kit (New England Biolabs). AtRecA1ΔC24 and AtRecA1ΔC28 were constructed by PCR using primers 19-20 and 21-22, respectively (**Table S1**). The AtRecXΔDUF construct was built using PCR from synthetic gene AtRecX (**pTwist vector, TableS1**) and cloned into bacterial expression vectors.

### Protein expression and purification

SSB1 (19.63 kDa), SSB2 (21.91 kDa), WHY1 (21.81 kDa), WHY2 (21.95 kDa), WHY3 (21.74 kDa), OSB1 (26.62 kDa), OSB2 (34.86 kDa), OSB3 (44.38 kDa), OSB4 (33.87 kDa), AtRadA (61.58 kDa) and MBP-AtRecG (148.11 kDa) proteins were expressed in BL21Solu-pRARE2 *E. coli* strain (**Table S2**). MBP-AtRecA1 (87.11 kDa), MBP-AtRecA2 (86.41 kDa), MBP-AtRecA3 (83.37 kDa), MBP-AtRecX (87.89 kDa) and their mutants were expressed either in BL21Solu, BL21Solu-pRARE2 or BL21Solu-pKJE7 *E. coli* strains (**Table S2**). All these proteins were expressed and purified as described in the supplementary methods. *E. coli* RecA and SSB were purchased from New England Biolabs and PROMEGA, respectively.

### DNA substrates preparations

#### cssDNA

The 3,276-nt circular ssDNA plasmid was produced using a starting culture of Briefly, 5 mL of 2x YT medium supplemented with tetracycline/ampicillin was inoculated with XL1-Blue strain carrying pGEM3Zf(+)-HOM vector and incubated at 37 °C for 12 h until saturation. The culture was diluted with fresh 2x YT medium containing VCSM13 helper bacteriophage (Agilent) at a concentration of 6×10^7^ pfu/mL and incubated at 37 °C for 1 h. To select the infected cells, kanamycin was added and the incubation continued for another 18 h. Supernatant was recovered by centrifugation, and phage particles were precipitated by adding a 20% PEG8000, 2.5 M NaCl solution on ice for 2 h. Pellet was recovered by centrifugation and resuspended in PBS buffer. Extraction was continued using E.Z.N.A.® M13 DNA Mini Kit (Omega BIO-TEK), following manufacturer instructions (29).

#### ldsDNA

dsDNA plasmid was producing from overnight culture inoculated with XL1-Blue strain carrying pGEM3Zf(+)-HOM vector using ZymoPURE™ II Plasmid Midiprep (Zymo Research) Kit, following manufacturer instructions. Linear double-stranded pGEM3Zf(+)-HOM was generated from a restriction digestion with *Kpn*I enzyme, for 2h at 37 °C. DNA was purified using QIAquick PCR purification spin columns (Qiagen) according to the protocol provided by the manufacturer.

#### Supercoiled plasmid

Supercoiled M13mp18 plasmid was extracted as described by Liu et al. 2011. Unless stated otherwise, DNA concentrations are expressed in moles of nucleotides.

### ATPase assay

ATPase activity was measure as previously reported (30) in reactions of 10 µL in 1X buffer (25 mM Tris-HCl pH 7.5, 1 mM DTT, 10 mM NaCl and 10 mM MgCl_2_) with 1 µL of a 1:1000 dilution radioactive γ^32^P-ATP (3300 Ci/mmol) and 1 mM of cold ATP. All reactions were carried out at 37 °C for 60 min. and terminated by addition of EDTA (80 mM). To determine the DNA substrate influence on the activity, 150 nM (8.5 μM nt) of either ssDNA (Oligo 1, **Table S3**) or 5’overhang (4.5 μM nt) (**Oligo 1 and 2, Table S3**) substrate was added with 150 nM of AtRecA protein. 3 µL aliquots of each reaction mixture was spotted on a polyethylene cellulose TLC. The reaction components [ATP and inorganic phosphate (Pi)] were separated by thin layer chromatography (TLC) using 0.4 M LiCl_2_ and 0.1mM EDTA as mobile phase. The dried TLC sheets were visualized by Storm 840 PhosphorImager (BioRad).

### D-loop assay

Briefly, 10 μL reactions were performed at 37 °C, with a reaction buffer containing 50 mM triethanolamine-HCl pH 7.5, 3mM DTT, 2 mM ATP, 1 mM MgCl_2_, 0.1 mg/mL BSA, 2 % glycerol, 12 mM phosphocreatine and 10 U/mL creatine kinase(31). The 5’ Cy5-labeled 90-mers oligo (187.5 µM nt; 2 nM in molecules) and the indicated concentration of either EcRecA or AtRecA were incubated for 10 min, to allow RecA filament formation. To initiate the formation of D-loop, the supercoiled plasmid M13mp18 RFI (14.41 µM nt; 1 nM in molecules) was added and MgCl_2_ concentration was increased to 10 mM and incubated for 8 min. The reactions were terminated by addition of 25 mM Tris-HCl pH 7.5, 2% SDS and 8 % glycerol. Products were analysed by electrophoresis in 0.8 % agarose gel and 1x TAE at 48 V for 70 min at room temperature (32). Gels were visualized on the Amersham Typhoon imager. Each experiment was repeated at least three times, and quantification was carried using ImageJ.

### Three-strand DNA exchange reaction

Unless stated otherwise, DNA concentrations are expressed in moles of nucleotides. All reaction incubations were carried out at 37 °C. 10 μM nt cssDNA pGEM3Zf(+)-HOM was incubated with 1X buffer (25 mM Tris-acetate pH 7.5, 3 mM DTT, 1 mM magnesium-acetate, 3 mM potassium glutamate, 0.1 mg/mL BSA and 5 % (w/v) glycerol) and 5 mM ATP, 10 U/mL creatine kinase and 12 mM phosphocreatine was used as ATP regeneration system (33,34). Unless otherwise indicated, 3.3 μM of either EcRecA or AtRecA was incubated for 10 min. Unless otherwise indicated, 2 μM of EcSSB or Arabidopsis SSBs were added, and the reaction was incubated for another 10 min. Each reaction was initiated by the addition of 20 μM nt ldsDNA pGEM3Zf(+)-HOM (*Kpn*I-linearized), and magnesium-acetate concentration was increased to 10 mM. After reactions were incubated for indicated times 10 μL aliquots were taken at appropriate times and added to 5 μL of 3:2 6X Ficoll:10 % SDS and further incubated for 10 min. Samples were subjected to electrophoresis on a 0.8 % agarose gel using TBE 1X, at 37 V for 12 h at room temperature. The gels were stained with ethidium bromide for 1 h and then washed two times with distilled water for 10 min. Gel images were captured by GelDoc Go Imaging System and DNA bands were quantified using ImageJ. Each experiment was repeated at least two or three times. For each reaction or lane, the 100 % of DNA represented the sum of the nicked circle (NC), join molecule (JM), ldsDNA and cssDNA. For strand exchange reactions with helicases AtRadA (250 nM as undecamer) or AtRecG (250 nM as monomer), were added to achieve a ratio of helicase:RecA 1:13. Standard conditions described above were followed at the RecA and SSBs protein concentrations, and helicases were added at different steps of the reactions as indicated in each experiment. In strand exchange reactions for inhibition activity assay with AtRecX, it was added at different steps of the reaction at the concentrations indicated in the figures.

### Recombination intermediate isolation and branch migration assays

Strand exchange reactions were scaled up to 450 μL, using AtRecA1 and WHY1 at the concentrations used in standard conditions described above. Reactions were stopped after 15 min by addition of EDTA to 12 μM, SDS to 0.7 %.and further incubation at 37 °C for 20 min (35,36). Reaction was deproteinized by phenol-chloroform extraction followed by ethanol precipitation. The DNA pellet was solubilized in buffer (25 mM Tris-acetate pH 7.5, 3 mM DTT, 10 mM magnesium-acetate, 3 mM potassium glutamate and 0.7 % (w/v) glycerol) and analyzed on a 0.8 % agarose gel, 1X TBE, at 75 V for 2.5 h at room temperature. DNA was used the same day for branch migration assays.

Branch migration assays were carried out at 37 °C with 1X buffer (25 mM Tris-acetate pH 7.5, 3 mM DTT, 10 mM magnesium-acetate, 3 mM potassium glutamate, 0.1 mg/mL BSA and 5 % (w/v) glycerol), 5 mM ATP, 10 U/mL creatine kinase and 12 mM phosphocreatine, and DNA intermediates. Reactions were initiated by the addition of the indicated amounts of AtRadA or AtRecG helicases or EcRecA. Aliquots of 10 μL were taken at the indicated times and added to 5 μL of 3:2 6X Ficoll:10 % SDS and further incubated for 15 min at 37 °C. Reactions were analyzed on a 0.8 % agarose gel, 1X TBE, at 37 V for 12 h at room temperature. The gels were stained with ethidium bromide for 1 h and then washed two times with distilled water for 10 min.

## RESULTS

### Plant organellar HR enzymes arise from diverse evolutionary origins

Plant mitochondrial and chloroplast HR and DNA replication systems harbor enzymes form diverse evolutionary origins (5,8,11,23,26). Several enzymes retain an unmistakable bacterial origin, while others are plant-specific innovations, nuclear-derived factors, or arisen from a T-odd bacteriophage lineage (**Table 1 and Table S1**). All HR plant organellar proteins are encoded in the nucleus and are targeted to mitochondria or chloroplast or have dual localization. For instance, *A. thaliana* SSB1and SSB2 are solely located into mitochondria (37,38), whereas members of the Whirlies, OSBs, and RecAs proteins can be targeted to one organelle or have dual localization. Some HR plant organellar proteins appear to have redundant or specialized functions and others like RecA2 and RecA1 are essential (39) (**Table 1**). Building on the experimental and genetic evidence supporting HR pathways proposed by José Gualberto’s group and others (5,6,11,22,26,28,36,40–43), we purified all components identified to date from the plant model *Arabidopsis thaliana* aiming to reconstitute and dissect plant organellar homologous recombination (**Table 1**).

**Table 1.**
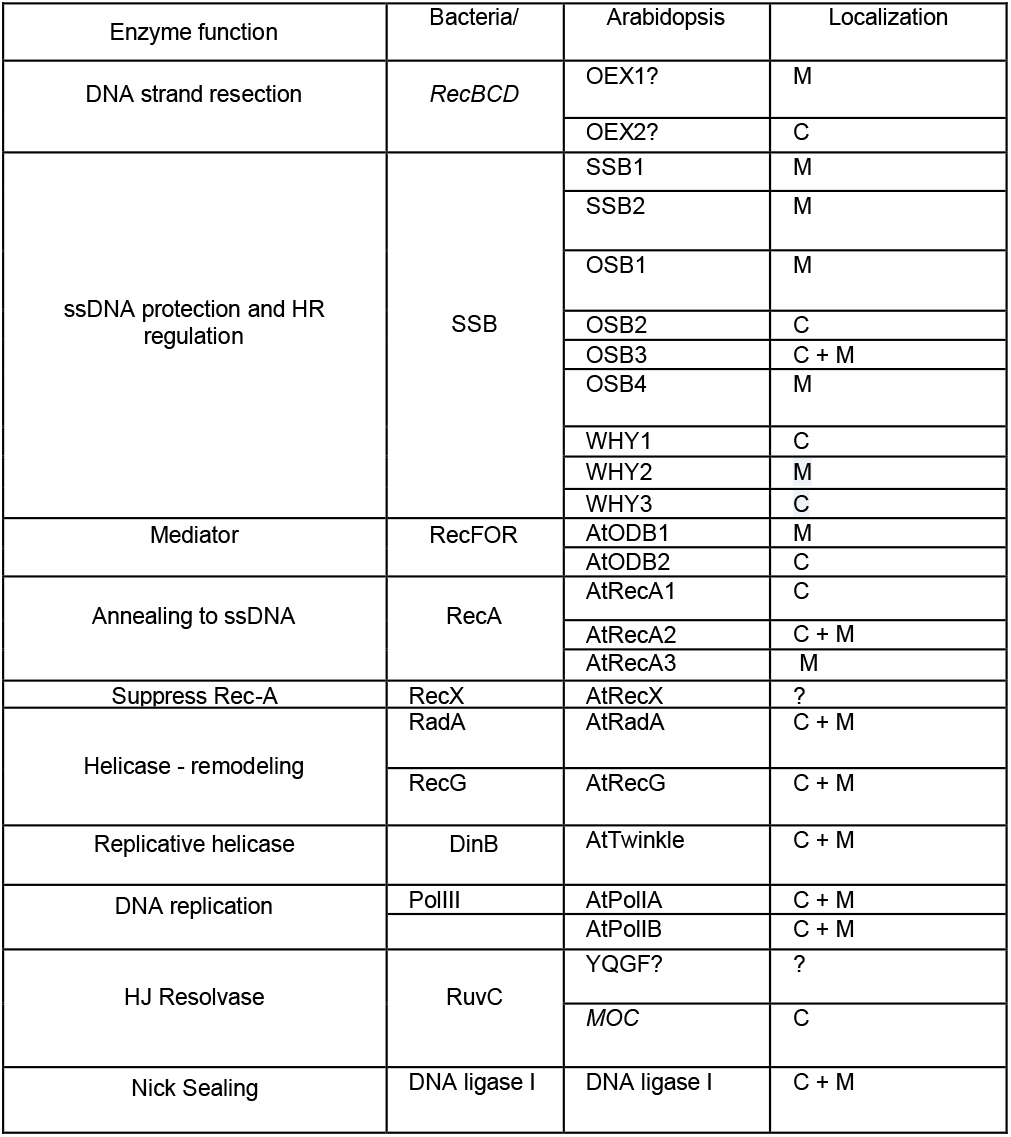
Minimal Set of HR Factors in Plant Organelles.

### Bacterially derived plant RecA proteins function as ATPases that promote D-loop formation but are inefficient at catalyzing extensive DNA strand exchange

In contrast to metazoan mitochondria, plant organelles retain RecA homologs of bacterial origin. *Arabidopsis* contains three bacterial-like RecA proteins with distinct organellar targeting: RecA1 is targeted to chloroplasts, RecA2 is targeted to both chloroplasts and mitochondria, and RecA3 is exclusively targeted to mitochondria (39,42,44). RecA1 is essential, RecA2 is required after seedling development, and RecA3 is dispensable under normal growth but important for resistance to genotoxic stress. Loss of RecA3 increases rearrangements at imperfect repeats, suggesting greater reliance on error-prone, RecA-independent pathways such as single-strand annealing. (42). Plant organellar RecA consist of a conserved central ATPase domain flanked by N- and C-terminal regions that may regulate filament dynamics and recombination efficiency (**Fig. 1A, 1B).** Plant RecAs conserved all motifs and amino acids involved in ATP hydrolysis, however RecA3 has a shorter C-terminal extension (**Fig. 1**). Plant RecA proteins were purified as MBP fusions, and assayed following tag cleavage (**Fig. 1D)**. In the absence of ssDNA, the ATPase activity of plant RecA proteins is barely detectable, whereas ssDNA regions stimulate ATP hydrolysis. Thus, like other RecA homologs, plant RecAs function as ssDNA-dependent ATPases (45,46) (**Fig. 1E).**

**Fig. 1.**
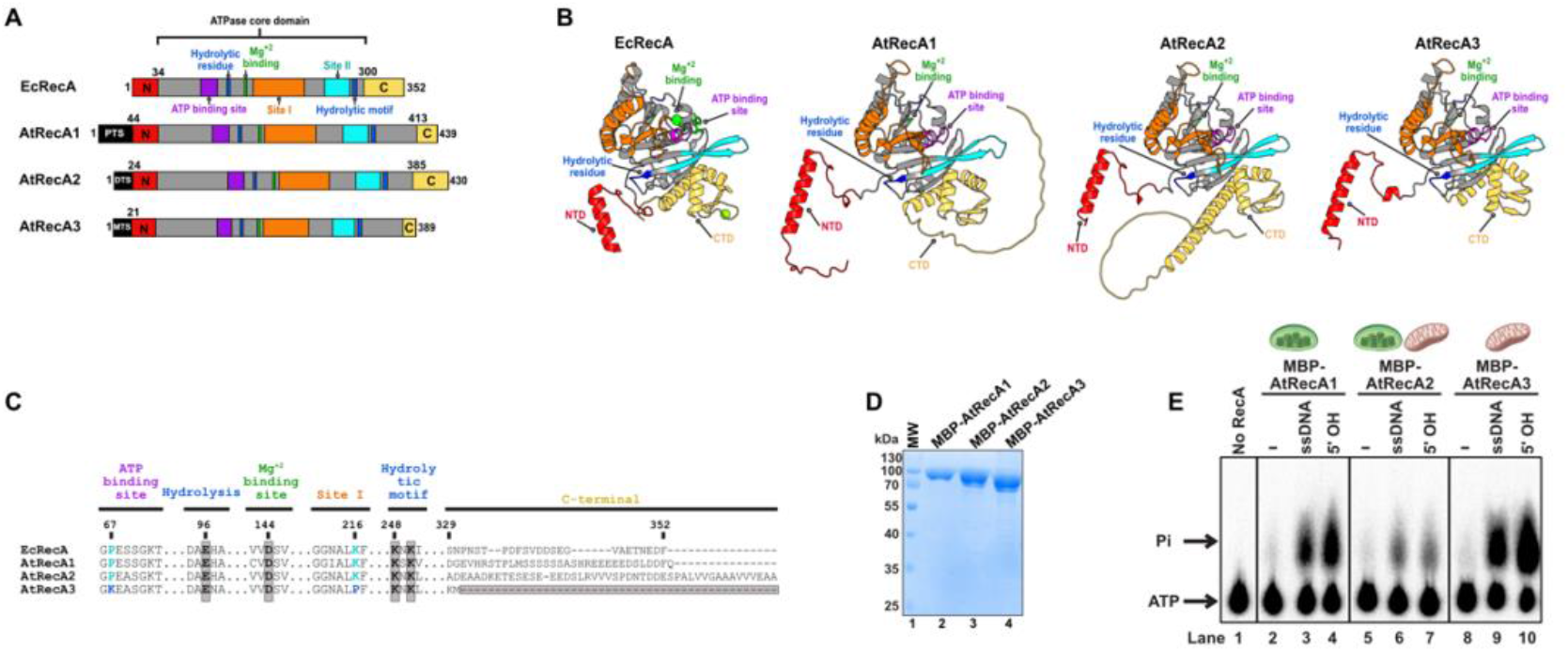
Plant organellar RecA proteins display DNA-dependent ATPase activity. A) Structural organization of AtRecA1, AtRecA2, and AtRecA3, corresponding to chloroplast-, dual-, and mitochondria-localized isoforms, respectively. AtRecA3 lacks the distinctive C-terminal extension present in AtRecA1 and AtRecA2. B) Structural homology models of AtRecA1, AtRecA2, and AtRecA3 in comparison to the crystal structure of *E. coli* RecA. C) Amino-acid sequence alignment of the three AtRecAs highlighting two key residue substitutions unique to AtRecA3: Lys463 in place of a conserved Pro, and Pro615 in place of a conserved Lys. D) 12% SDS–PAGE of showing the recombinant purification of full-length AtRecA proteins fused to MBP lacking their organellar targeting peptides. E) TLC assay showing ATP hydrolysis by AtRecAs in the presence (single-stranded oligonucleotide of 57 nt (ssDNA) and dsDNA harboring a 5’-overhang (5’OH) of 30 nts) and absence of DNA (-). Arrows indicate initial labeled substrate ATP and inorganic phosphate (Pi) as the labeled released product.

Recombinases initiate homologous recombination by pairing ssDNA with a complementary strand and displacing the original strand to assembly a displacement loop (D-loop) **(Fig. 2A)**. D-loop formation represents the earliest step of homologous recombination and provides a direct readout of RecA-mediated homology recognition and strand invasion, making it a fundamental assay for dissecting RecA recombinase activity. To assess the recombinase activity of AtRecA proteins, we performed D-loop formation assays that measure ATP-dependent homologous pairing and strand exchange. In this experiment, a 90-nt FAM-labeled oligonucleotide was preincubated with increasing concentrations of AtRecA proteins and *E. coli* RecA as positive control, followed by the addition of homologous supercoiled plasmid DNA. All AtRecA proteins promoted the formation of D-loop intermediates, confirming their ability to catalyze homologous recombination reactions. AtRecA1 exhibited substrate-dependent D-loop formation comparable to that of *E. coli* RecA, AtRecA2 efficiently promotes D-loop formation, but increasing concentrations of AtRecA2 reduce D-loop accumulation, suggesting D-loop dissociation or unwinding, as reported for *E. coli* RecA (47)(**Fig. 2B** and **2C**). AtRecA3 displayed reduced efficiency in D-loop formation in comparison to AtRecA1 and AtRecA2 (**Fig. 2B and 2C**). Albeit D-loop formation is commonly used to assess HR, the sequence of events underlying three-strand HR is fully captured in the strand-exchange reaction between homologous circular single-stranded DNA (cssDNA) and linear double-stranded DNA (ldsDNA). In this experiment, a recombinase polymerizes on ssDNA to form a presynaptic filament, that engages homologous linear dsDNA to establish synapsis and initiate heteroduplex formation between the cssDNA circle and the complementary strand of the duplex. As strand exchange progresses, one strand of the linear duplex is progressively displaced while the remaining strand becomes fully paired with the circular ssDNA, resulting in a nicked circular heteroduplex DNA (NC) and a displaced linear single-stranded DNA molecule (**Fig. 2D**). Branch migration mediated by *E. coli* RecA occurs at a slow rate and accordingly, RecA-catalyzed strand-exchange reactions are typically incubated for extended periods, often several hours, to allow accumulation of joint molecules (JM) and the subsequent formation of final exchange products (32,48). In control reactions, *E. coli* RecA promotes complete strand exchange between linear duplex DNA and cssDNA, yielding nicked circular products on approximately 41.7% of the substrate within 120 min in the absence of EcSSB (**Fig. 2E, lane 5**). In contrast, the AtRecA proteins in the absence of any plant SSBs, are largely impaired in the completion of strand exchange, leading primarily to the accumulation of joint-molecule intermediates (**Fig. 2E, lane 7 to 39**). Only AtRecA1 generates nicked circular products, which are detected after prolonged incubation times (180–240 min) and reach a maximum yield of less than 10% (1.7 % and 6.9 %, respectively) of the input substrate (**Fig. 2E**, **Fig. 2F**). The inability of AtRecAs to complete strand exchange resemble other recombinases, such as bacteriophage T4 UvsX which accumulate joint-molecule intermediates and require accessory helicases, such as UvsW, to complete strand exchange (34,49–52).

**Fig. 2.**
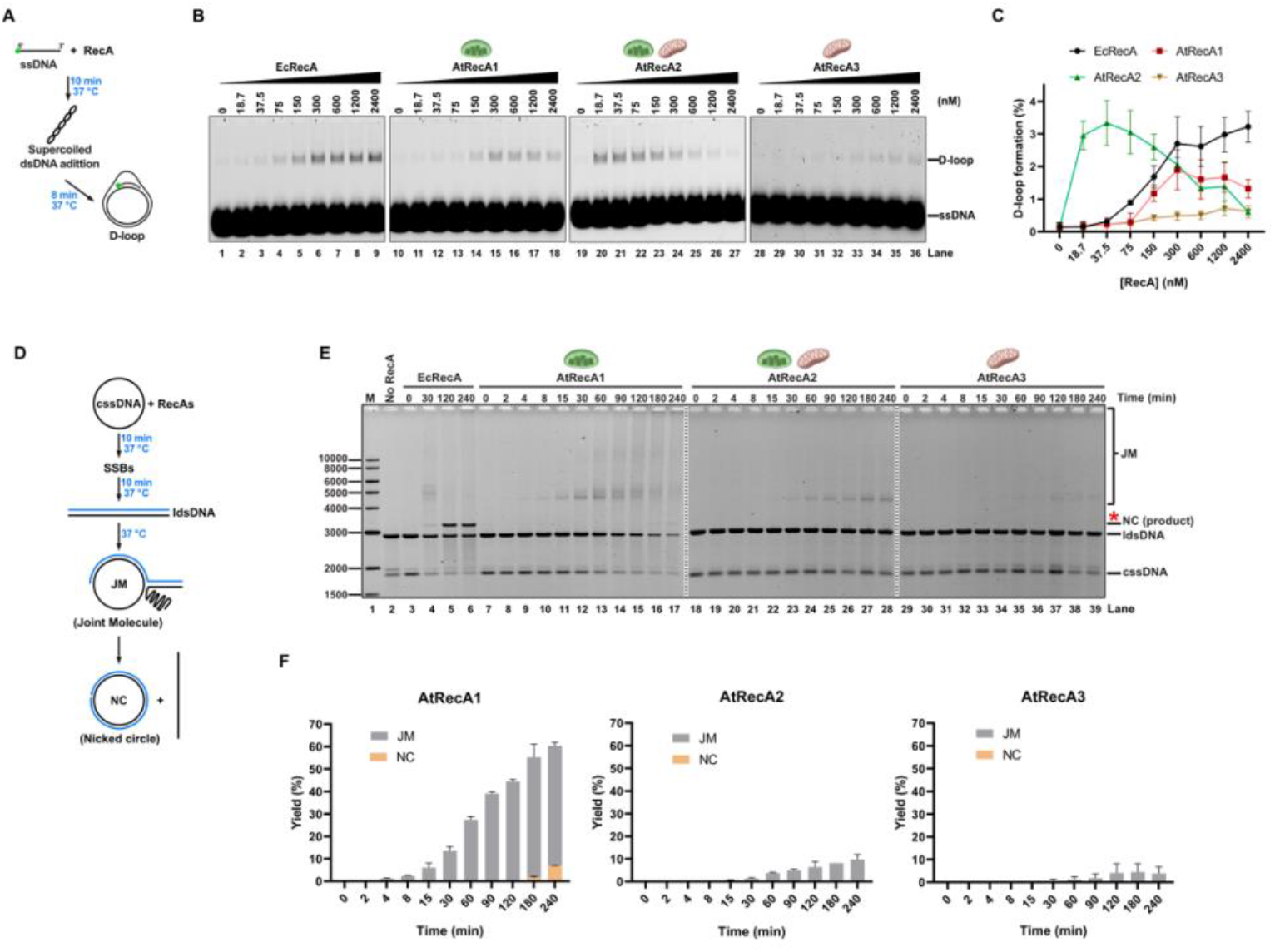
Plant organellar RecAs catalyze D-loop formation and exhibit differential strand-exchange activities without SSB protein. **A)** Schematic representation of D-loop formation mediated by AtRecA proteins. **B)** Native agarose gel showing the migration and the quantification of a Cy5-labeled 90-nt oligonucleotide homologous to supercoiled M13mp18 plasmid after an incubation period of 8 minutes, and increased concentrations of RecAs, visualized by fluorescence scanning. **C)** Quantification of D-loop formation by AtRecAs. Each experiment was repeated at least three times, and quantification was carried using ImageJ. **D)** Schematic representation of three-strand DNA exchange reactions catalyzed by RecAs. Circular ssDNA (cssDNA), linear dsDNA (ldsDNA), joint molecule (JM) intermediates and nicked-circle (NC) end-product. **E)** Strand exchange reaction time courses catalyzed by RecAs without plant SSB proteins. 10 μM nt cssDNA pGEM-HOM was preincubated with 3.3 μM of AtRecA or EcRecA for 10 min. For initiating reaction 20 μM nt ldsDNA pGEM-HOM was added, and magnesium-acetate concentration was increased to 10 mM. Aliquots of 10 μL were taken at appropriate times and stopped. Samples were subjected to electrophoresis on a 0.8 % native agarose gel, stained with ethidium bromide showing the formation of joint-molecule intermediates in reactions containing AtRecA proteins in comparison to RecA from *E. coli*. AtRecA2 and AtRecA3 fail to generate detectable complete nicked circular final product and only joint molecule intermediates were observed. AtRecA1 produces nicked circle at the longest times, the characteristic nicked-circular product of homologous recombination as is shown with EcRecA as positive control. **F)** Quantification of nicked-circle end-product and joint molecule intermediates formation by AtRecAs. Data in C and F represents the mean values +/− standard deviation from triplicate experiments.

### Plant unique ssDNA-binding proteins potentiate DNA strand-exchange by AtRecAs

SSB proteins are traditionally viewed as factors that protect ssDNA from exonucleolytic degradation and must be displaced by recombination mediators to facilitate RecA nucleation and filament formation during homologous recombination (53,54). In contrast, to this notion, bacterial SSB stimulates RecA filament formation (55). Plant organelles contain a large and functionally diverse repertoire of SSB proteins, comprising nine members distributed in three structurally and evolutionary distinct protein families. Plant organellar SSBs include two bacteria-like SSBs, homologous to metazoan mitochondrial SSBs and exclusively targeted to mitochondria, which we refer to herein as mtSSB1 and mtSSB2 (25,37). In addition, plants possess two plant-specific families of SSB proteins: four organellar single-stranded DNA-binding proteins (that form the OSBs family), targeted to mitochondria and plastids, and three Whirly (WHY-family) proteins (Reviewed in (5,22,27,28,56)) (**Fig. 3A and Table 1**). Cellular biology studies indicate that plant SSBs modulate DNA repair, for instance WHY2 and WHY3 impinge MMEJ (57), OSB1 and OSB4 regulate ectopic recombination across intermediate-sized repeats (43), and mtSSB1 and mtSSB2 participate in homologous recombination (37,38). Although mtSSBs, OSBs, and Whirlies have been implicated in HR, their direct roles have not been tested. Here, we directly test their roles in HR using purified proteins of all nine organellar SSBs (**Fig. 3B**). The plastid-localized AtRecA1 and mitochondria-localized AtRecA3 were tested exclusively with plastid- and mitochondrial-localized SSBs, respectively, whereas the dual-targeted AtRecA2 was tested with all nine organellar SSB proteins. Nicked circular product formation was evaluated relative to the stimulation observed for *E. coli* RecA in the presence of *E. coli* SSB (**Fig. 3C, E, G, lanes 2 to 3**). When AtRecA1 was incubated with the chloroplast ssDNA-binding proteins WHY1, WHY3, OSB2 or OSB3, JM intermediates accumulated rapidly and were detectable within 2 min (**Fig. 3C)**. The chloroplast-localized SSBs, WHY1, WHY3, OSB2, but not OSB3 stimulated AtRecA1-mediated strand exchange, resulting in the formation of nicked circular products in comparable amounts to the *E. coli* RecA control (**Fig. 3C and 3D**). For instance, the addition of WHY1 and WHY3 result in 41 % and 51% of NC final product in comparison to less than 10% in their absence (**Fig. 3C, lanes 5 to 26)**. Reactions containing AtRecA1 and OSB2 produced NC (15 %), and DNA aggregates (83 %) that remained trapped in the loading wells. These species likely arise from repeated intramolecular strand invasion and synapsis events, leading to the formation of complex branched DNA structures, reminiscent of the branched DNA networks reported for bacteriophage T4 UvsX-mediated recombination reactions (52,58) **(Fig. 3C, lanes 27 to 37)**. For AtRecA2, the addition of SSBs have minimal effect of the formation of the nicked-circle final product, as only reactions incubated with Whirlies promote less than 5% of NC product **(Fig. 3F)**. However, the plant-specific SSB proteins WHY1, WHY2 and WHY3 increased the formation of intermediate joint-molecule products (48 %, 60 % and 41 %, respectively) compared to reactions without plant SSBs (9.5 % of JM) within the maximum reaction time **(Fig. 3E, lanes 27–59)**. By contrast, mtSSB1, mtSSB2, OSB1, OSB3, and OSB4 reduced the accumulation of joint-molecule intermediates in reactions containing AtRecA2, whereas OSB2 did not affected joint-molecule formation (12 %). For AtRecA3, solely WHY2 and OSB3 produced a slight increase in the appearance of join molecules (2.3 % and 8 %, respectively), as reactions incubated with AtRecA3 in absence of SSBs only accumulate a barely visible amount of joint molecule products (1.5 %). Regarding the formation of the nicked-circle final product, only WHY2 has minimal stimulatory effect (3.2 %) **(Fig. 3G and 3H, lanes 1 to 70**). Our data shows that plant whirlies proteins but not canonical mtSSB1 and mtSSB2 or OSB2 promote complete strand exchange by AtRecA1, in contrasts in reactions catalyzed by AtRecA2, Whirlies and OSBs enhance the formation of joint molecules but not NC and in reactions incubated with AtRecA3 only reaction incubated with WHY2 produced NC products.

**Fig. 3.**
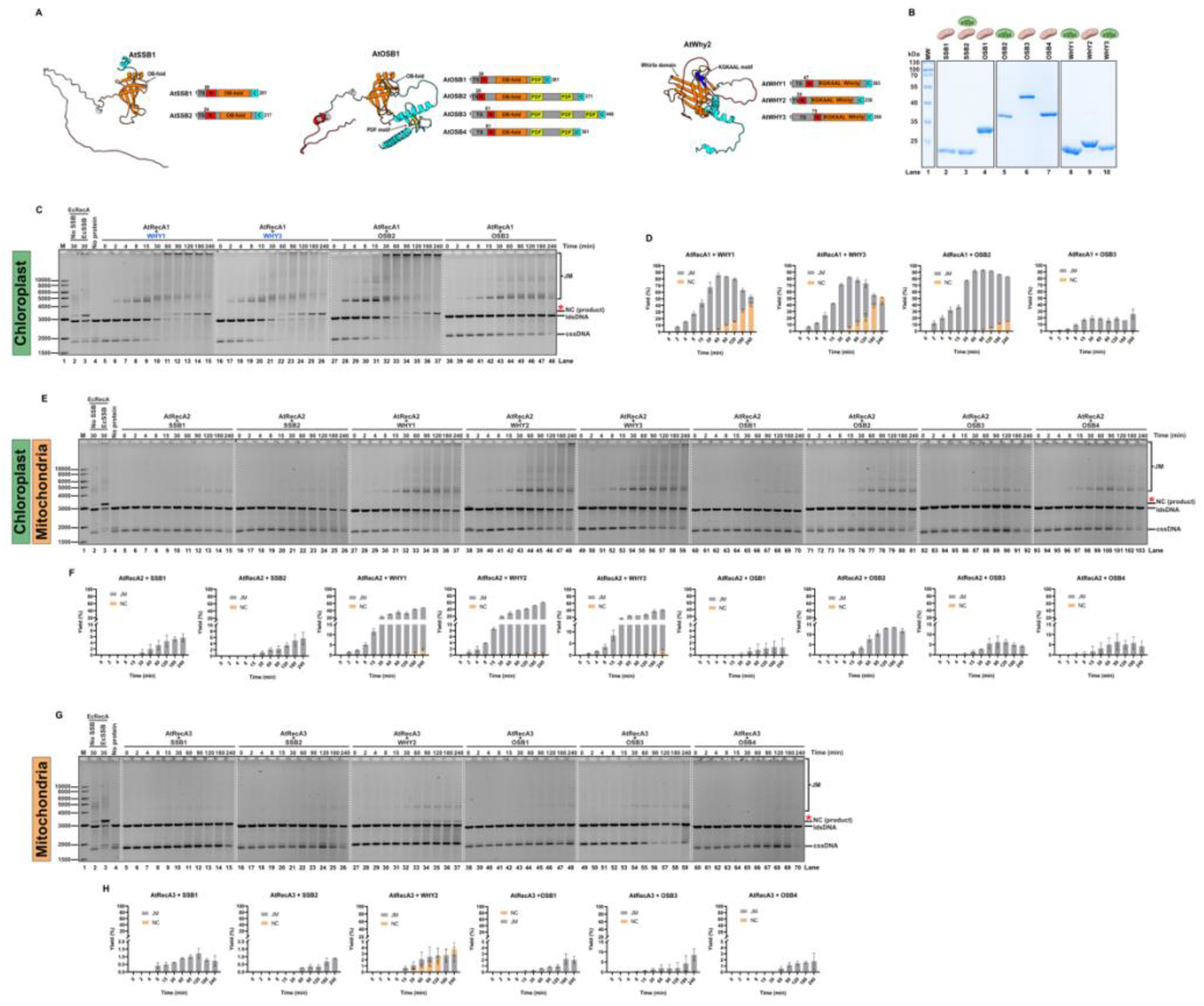
Plant organellar ssDNA-binding proteins potentiate HR. **A)** Structural homology models and graphical representations of the canonical AtSSBs (SSB1 and SSB2), the plant-specific Organellar Single-Stranded DNA-Binding proteins (OSBs), and the Whirly family (WHY), illustrating the diversity of ssDNA-binding proteins (SSBs) architectures within plant organelles. **B)** 15% SDS–PAGE showing the heterologous purification of all nine plant organellar SSBs. **C, E, G**) Time-course strand-exchange reactions catalyzed by AtRecAs in the presence of their cognate organellar SSBs. 10 μM nt cssDNA pGEM-HOM was preincubated with 3.3 μM of AtRecAs or EcRecA for 10 min. 2 μM of EcSSB or Arabidopsis SSBs were added, and the reaction was incubated for another 10 min. For initiating reaction 20 μM nt ldsDNA pGEM-HOM was added, and magnesium-acetate concentration was increased to 10 mM. Aliquots of 10 μL were taken at appropriate times and stopped. Samples were subjected to electrophoresis on a 0.8 % native agarose gel, visualized by ethidium bromide-stained showing the formation of strand-exchange intermediates and final end-products**. C and D)** WHY1 and WHY3 stimulated strand exchange by AtRecA1, leading to the appearance of NC products after 60 min and complete substrate conversion by 240 min. In contrast, OSB proteins supported only limited NC product formation. Reactions containing OSB2 produced high-molecular-weight DNA aggregates that remained trapped in the loading wells, consistent with the formation of complex branched DNA networks. **E and F)** AtRecA2 NC product formation is minimally potentiated by WHY1, WHY2 and WHY3 (lanes 27 to 59), but the accumulation of JM intermediates is enhanced, whereas mtSSB1, mtSSB2, OSB1, OSB3 and OSB4 reduced JM formation, and OSB2 had little effect on AtRecA2 activity. **G and H)** AtRecA3 displayed limited strand-exchange activity solely in the presence of WHY2, whereas the addition of OSB3 resulted in JM accumulation. Data presented in D, F, and H represents the mean values +/− standard deviation from triplicate experiments.

### Plant DNA helicases AtRadA and AtRecG catalyze branch migration on isolated intermediate joint molecules

AtRecAs catalyze a substantial amount of JM intermediates and aggregates due to its limited intrinsic ability to promote branch migration during strand exchange reactions (**Figs. 3C, 3E and 3G**), in bacteriophage T4, the helicase UvsW promotes the processing of recombination intermediates. These observations raise the possibility that plant helicases may likewise play an important role to promote the formation of nicked circular final product from branched DNA intermediates. Plant organelles rely on at least two DNA helicases for homologous recombination: AtRadA and AtRecG (26,36,59) (**Fig. 4A**). The proposed function of AtRadA parallels the RadA/RecA interplay in *Bacillus subtilis* and *Streptococcus pneumoniae,* in which RadA acts downstream of RecA to promote bidirectional unwinding by loading onto ssDNA and driving dsDNA unwinding and rewinding to facilitate heteroduplex extension (36,60,61). In contrast, the role of the DNA translocase RecG is not well-defined. RecG is a member of the SF2 helicase superfamily and an analog of bacteriophage T4 UvsW. RecG preferentially processes branched DNA substrates, including Holliday junctions, and stalled replication forks, through 3′–5′ directional DNA unwinding (62–65). To examine if AtRadA and AtRecG catalyze branch migration in the absence of RecA, we employed join molecule intermediates synthetized by a reaction incubated with AtRecA1 and WHY1 and phenol-chloroform purified to remove proteins. This DNA was then incubated with AtRadA or AtRecG in the presence of ATP to assess if the helicase alone could transform the intermediates into final products (**Fig. 4B and 4C**). When EcRecA was added to these intermediates, nicked-circle products were readily observed (65) (**Fig. 4D, lanes 2-4**). Without the addition of RecA or DNA helicases there is no progression of the reaction (**Fig. 4, lanes 5-7**). Addition of AtRadA and AtRecG, results in the resolution of branched recombination intermediates, leading to the accumulation of the final nicked double-stranded circular product (**Fig. 4D, lanes 8-13**). Collectively, these assays demonstrate that AtRadA and AtRecG act directly on purified recombination intermediates to promote their efficient transform into a nicked circular heteroduplex DNA molecule and a displaced linear single-stranded DNA as the final reaction products.

**Fig. 4.**
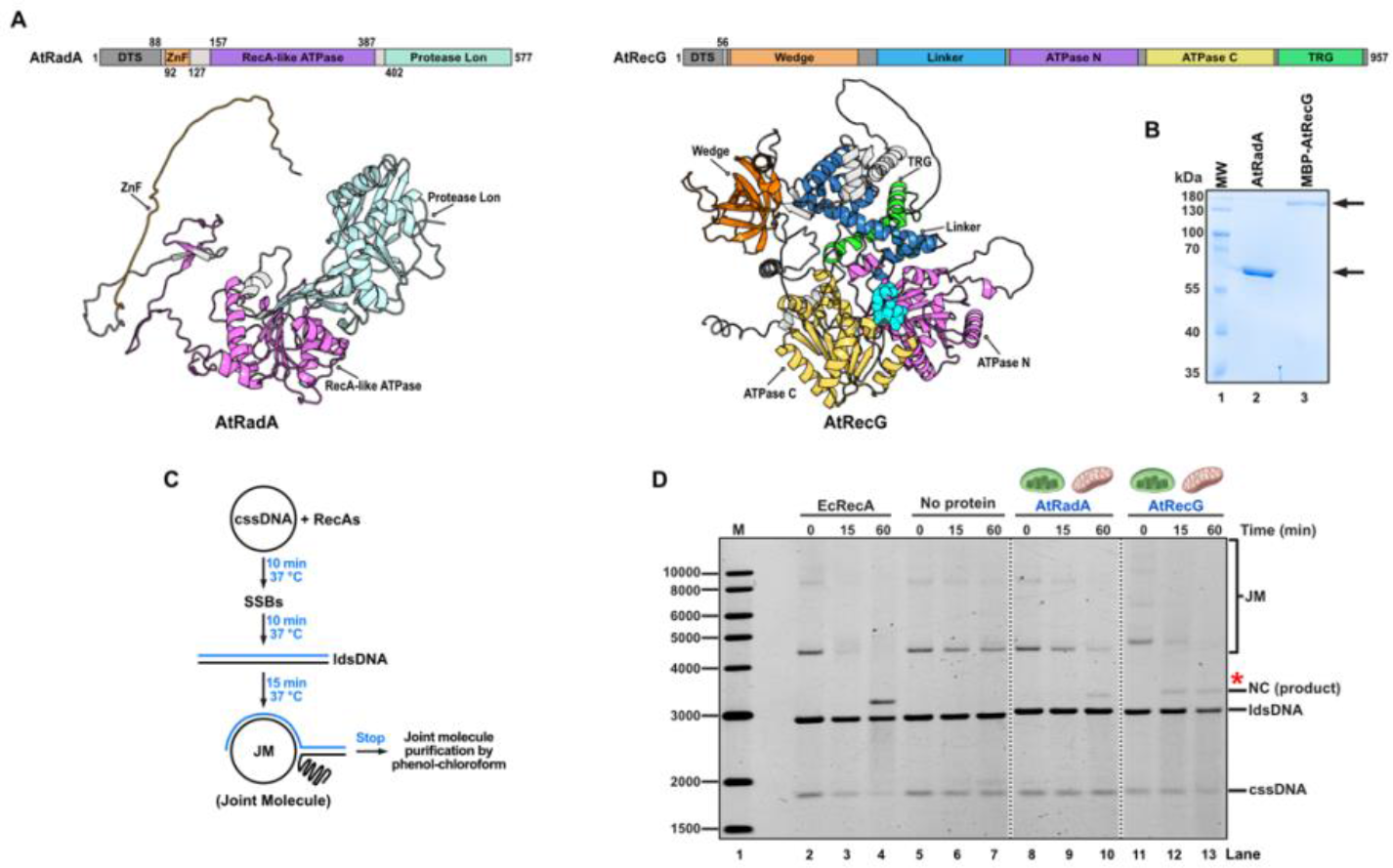
Plant organellar DNA helicases promote branch migration of preassembled RecA recombination intermediates. **A)** Domain organization of the plant organellar DNA helicases used in this study. AtRadA contains an N-terminal zinc-finger (ZnF) domain, a RecA-like ATPase domain, and a C-terminal Lon protease-like domain. AtRecG is composed of an N-terminal wedge domain, a linker region, two ATPase domains, and a C-terminal translocation in RecG (TRG) motif. **B)** 10% SDS-PAGE Coomassie-stained gel showing the purified recombinant proteins AtRadA (61.58 kDa) and MBP-AtRecG (148.11 kDa), following three purification steps. **C)** Schematic representation of joint-molecule (JM) intermediates generation by AtRecA1-mediated strand exchange and then phenol-chloroform-purified for the branch migration assay. **D)** Branch migration of purified JM intermediates, by AtRadA and AtRecG. Preassembled JM intermediates were purified and subsequently incubated with the indicated helicases or EcRecA, as indicated in methods. Addition of AtRadA and AtRecG promoted the branch migration of recombination intermediates, resulting in the accumulation of nicked circular (NC) DNA products.

### Plant DNA helicase AtRadA but no AtRecG facilitates AtRecA mediated DNA strand exchange by branch migration activity

To assess how coordinated protein activities influence the completion of strand-exchange reactions, we reconstituted homologous recombination reactions containing RecAs, SSBs, and DNA helicases. First, we assessed four different helicase-addition schemes to examine if the order of addition influences the apparition of final products (**Fig. S1 and Fig. S2**). In all cases, standard conditions were followed with circular ssDNA (cssDNA) preincubation with AtRecA, and the following addition schemes were tested: (A) helicase was added simultaneously with AtRecA, (B) helicase was added simultaneously with SSB, (C) helicase was added simultaneously with ldsDNA, and (D) helicase was added 15 min after reaction was in progress (**Fig. S1 and Fig. S2**). We found that the order of helicase addition has no significant effect on the formation of the final NC product in reactions catalyzed by AtRecA1 and AtRecA3, but in reactions with AtRecA2, there was a slightly increase on the yield of the NC product formation if a helicase was added simultaneously with ldsDNA (reaction order C), and for this reason in subsequent experiments we employed this order of addition (**Fig. S1 and Fig. S2**).

Next, we evaluated full-length product formation by AtRecAs over 240 min of incubation to determine whether AtRadA potentiates the conversion of intermediates into the final product (**Fig. 5**). In reactions containing the chloroplast-targeted recombinase AtRecA1 and AtRadA in the absence of SSBs, joint-molecule intermediates decreased over time, accompanied by the appearance of nicked-circle products beginning at 30 min. The final product reached a maximum yield of 13.7% after 240 min (**Fig. 5A and 5B, lanes 9 to13**). In same conditions, but adding chloroplast targeted WHY1, WHY3, and OSB2 the amount of joint molecules intermediates decreased (−24 %, −26 % and −50 %, respectively compared in reactions incubated in the absence of AtRadA) and the amount of nicked circle products increased (15 %, 16 % and 27 %) (**Fig. 5A and 5B, lanes 14 to 28**) (**Table S4**). Notably, reactions incubated with WHY1, WHY3, and OSB2 achieved nearly complete conversion to the nicked circular final product (57 %, 67 %, and 42 %, respectively) (**Fig. 5A and 5B, lanes 18, 23 and 28**). Reactions incubated with the dual targeted OSB3 showed an interesting effect, both JM and NC products increased (13.9 % and 18.5 %, respectively compared to reactions incubated without AtRadA), but the maximum yield of NC at 240 min was low (14.7 %) (**Fig. 5A and 5B, lanes 29 to 33**). These data demonstrate that although AtRecA1 and AtRadA are sufficient for final product formation in chloroplast-targeted reactions, high product yields depend on chloroplast-specific SSBs, particularly WHY1, WHY3, or OSB2.

**Fig. 5.**
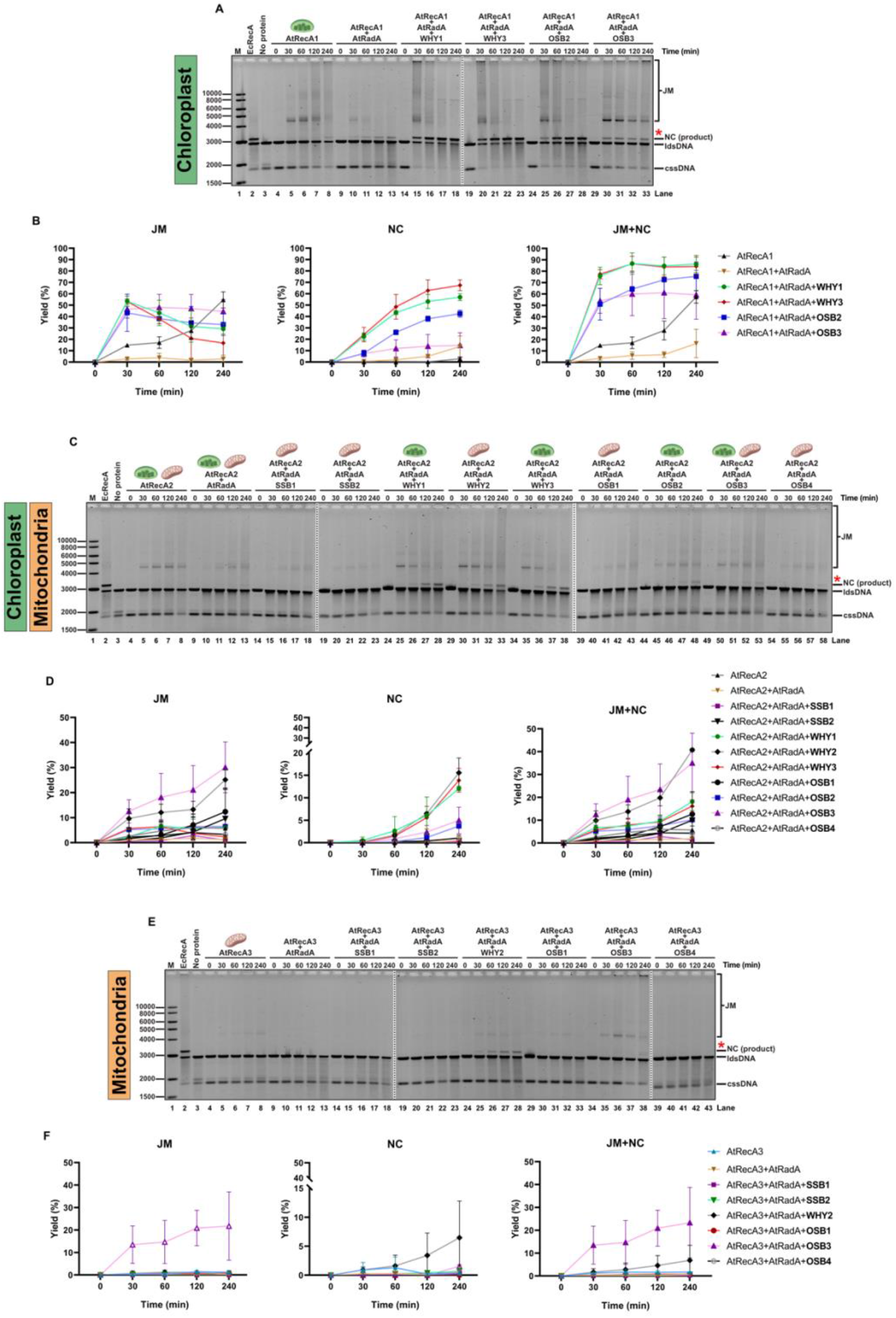
AtRadA facilitates the completion of RecA-mediated three-strand exchange reaction by branch migration activity. Time course reactions to evaluated AtRadA helicase effect on three-strand exchange reactions catalyzed by **A-B)** chloroplast targeted AtRecA1, **C-D**) dual targeted AtRecA2 and **E-F**) mitochondrial targeted AtRecA3, with their cognate SSB proteins. Reactions were assembled as described in methods: 10 μM nt cssDNA pGEM-HOM was preincubated with 3.3 μM AtRecA for 10 min, 2 μM SSB proteins were added and incubation continued for 10min. For initiating reaction 20 μM nt ldsDNA pGEM-HOM was added with 250 nM AtRadA. Aliquots were removed at the indicated time points and run on 0.8 % agarose gel. Gel staining was followed as described in methods. (cssDNA) circular ssDNA, (ldsDNA) linear dsDNA, (NC) nicked-circle final product and (JM) joint molecules intermediate product. **B, D and F**) The amounts of NC and JM products are indicated as the percentage of total substrate added.

In the absence of SSBs, reactions mediated by AtRecA2 and AtRadA produced detectable nicked-circle (NC) products only after 240 min, reaching a maximum yield of 1.1%. (**Fig. 5C, lanes 9 to13**). In reactions containing mtSSB1 or mtSSB2, joint-molecule levels decreased by 4.6% and 4.5%, respectively, relative to reactions without AtRadA. However, neither protein potentiated nicked-circle product formation (**Table S5, Fig. 5C, lanes 14 to 23**). In the presence of chloroplast targeted WHY1, WHY3 and mitochondrial targeted WHY2 joint molecules decreased (−42.3, −36.7 and −35.2 %, respectively compared without AtRadA) and NC formation have their maximum yields of 12 %, 13.8 % and 15.5 %, respectively (**Table S5, Fig. 5C and 5D, lanes 24 to 38**). Reactions containing chloroplast-targeted OSB2 or mitochondria-targeted OSB4 showed modest decreases in joint-molecule levels (5.4% and 0.7%, respectively) relative to reactions without AtRadA; concomitantly, nicked-circle product formation increased only modestly, by 3.7% and 1.0%, respectively. In contrast to the reactions described above, in which AtRadA decreased JM levels and increased NC formation, reactions containing mitochondria-targeted OSB1 or dual-targeted OSB3 showed increased JM levels (11.6% and 25.8%, respectively, relative to reactions without AtRadA). Consistent with this, NC product formation increased only modestly, by 0.6% and 4.9%, respectively (**Table S5, Fig. 5C and 5D, lanes 39 to 58**). Our data suggests that effective DNA strand exchange in homologous recombination reactions by AtRecA2 requires the AtRadA and WHY1 or WHY3 in chloroplast, and WHY2 in mitochondria. The ability of distinct chloroplast and mitochondrial ssDNA-binding proteins to promote AtRecA2-dependent recombination suggests that this dual-targeted recombinase has been integrated into organelle-specific HR pathways.

In reactions in which homologous pairing and strand invasion were initiated by AtRecA3, the addition of AtRadA was not sufficient to promote JM and NC products (**Fig. 5E, lanes 9 to 13**). In the same reactions supplemented with mitochondria-targeted SSB1, SSB2, OSB1, or OSB4, JM formation decreased, but no detectable NC product was observed, indicating that these proteins did not promote efficient conversion to the final product (**Fig. 5E and 5F**). In reactions containing mitochondria-targeted WHY2, NC product formation increased by 3% relative to reactions without AtRadA (**Table S6**), reaching a maximum yield of 6.4% (**Fig. 5E and 5F, lanes 24 to 28**). However, in reactions containing dual-targeted OSB3, JM formation increased by 14.7% relative to reactions without AtRadA (**Table S6**), while the final NC product was detectable, reaching 1.5%. (**Fig. 5E and 5F, lanes 34 to 38**). These results suggest the existence of a functional AtRecA3-dependent homologous recombination pathway in mitochondria, in which WHY2 acts as an accessory factor, and AtRadA helicase increases completion of homologous recombination final product.

AtRecG likely functions as a plant organellar counterpart of bacteriophage T4 UvsW, promoting branch migration and remodeling recombination intermediates. While genetic studies indicate that AtRecG plays a minor role in chloroplast double-strand break repair, it is essential for mitochondrial genome stability. Time course reactions with AtRecA1 and AtRecG in the absence of SSBs (**Fig. 6A and 6B, lanes 9 to13**), shows that the AtRecG helicase does not have a significant effect on NC formation (**table S4**). Reactions containing WHY1 or WHY3 (**Fig. 6A, B, lanes 14–23**) showed a notable increase in JM formation (19% and 18%, respectively, relative to reactions without AtRecG). In parallel, NC product formation decreased by 18%, reaching maximum yields of 23% and 33%, respectively—approximately half of the NC product obtained in the presence of AtRadA (**Table S4**). Reactions incubated with OSB2 (Fig. 6A,B, lanes 24–28) showed no significant change in NC formation relative to reactions incubated in the absence of AtRecG, although JM formation increased by 40% compared with reactions incubated with AtRadA (**Table S4**), whereas reactions incubated with dual-targeted OSB3 (**Fig. 6A, B, lanes 29–33)** showed a 32.3% increase in JM formation relative to reactions without AtRecG but produced only 4.8% NC product. Our data indicates that in chloroplast, AtRecG promotes an increase in branched intermediates and a significant decrease in the final product, indicating that AtRecG does not compensate for AtRadA.

**Fig. 6.**
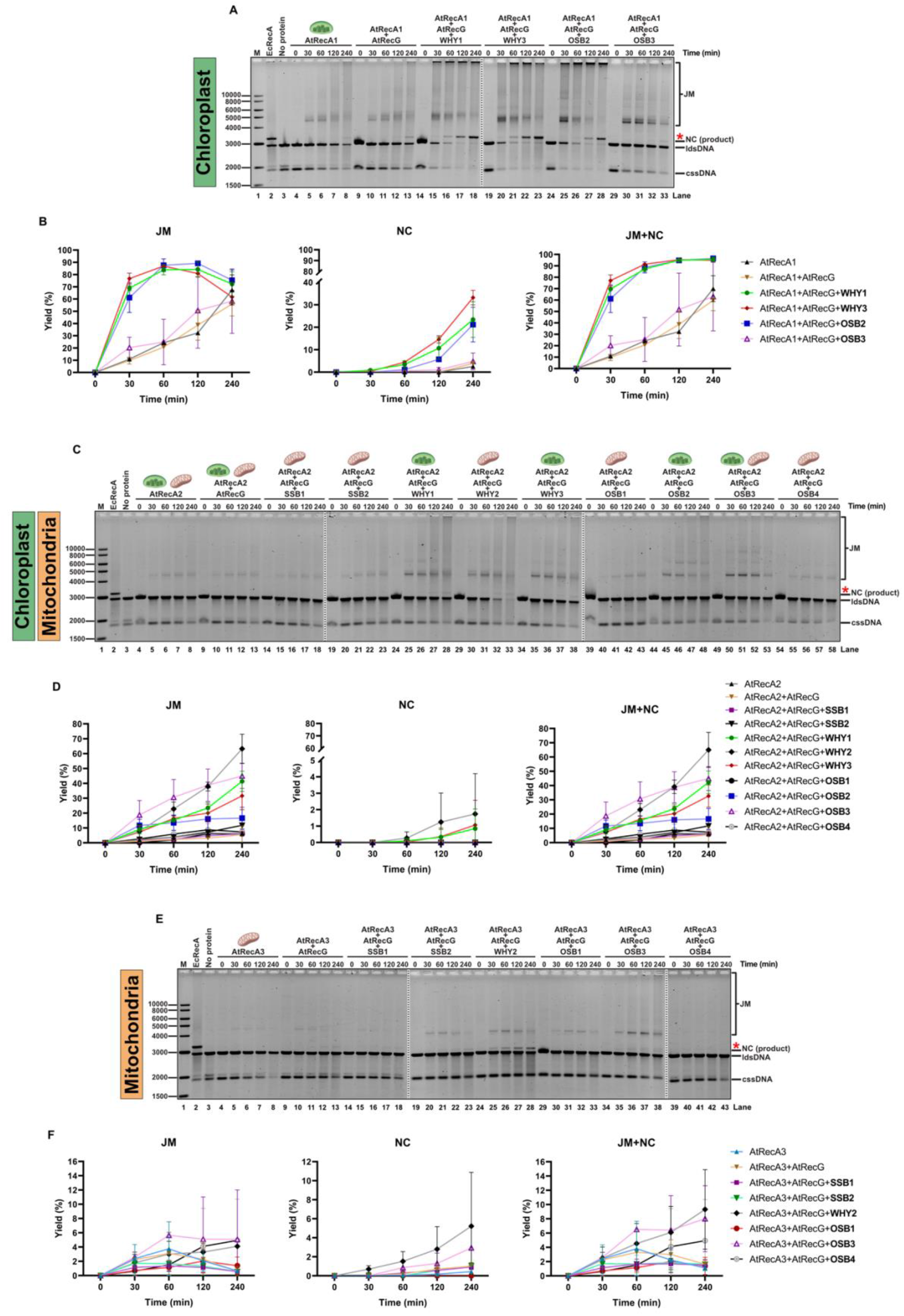
AtRecG promotes JM accumulation in three-strand exchange reaction mediated by AtRecA1 and AtRecA2, but not with AtRecA3. Time course reactions to evaluated AtRecG helicase effect on three-strand exchange reactions catalyzed by **A-B)** chloroplast targeted AtRecA1, **C-D**) dual targeted AtRecA2 and **E-F**) mitochondrial targeted AtRecA3, with their cognate SSB proteins. Reactions were assembled as described in methods: 10 μM nt cssDNA pGEM-HOM was preincubated with 3.3 μM AtRecA for 10 min, 2 μM SSB proteins were added and incubation continued for 10min. For initiating reaction 20 μM nt ldsDNA pGEM-HOM was added with 250 nM AtRecG. Aliquots were removed at the indicated time points and run on 0.8 % agarose gel. Gel staining was followed as described in methods. (cssDNA) circular ssDNA, (ldsDNA) linear dsDNA, (NC) nicked-circle final product and (JM) joint molecules intermediate product. **B, D and F**) The amounts of NC and JM products are indicated as the percentage of total substrate added

in reactions containing AtRecA2, SSBs, and AtRecG, the general trend was an increase in JM formation (Fig. 6C, D), while NC products were detected only with WHY1, WHY2, and WHY3 (Fig. 6C, D, lanes 24–38), with yields equal to or lower than those obtained in the absence of AtRecG (**Table S5**). Notably, reactions containing OSB3 (**Fig. 6C, lanes 49–53**) showed the greatest increase in JM formation (40.6% relative to reactions without AtRecG; **Table S5**). Thus, AtRecG does not promote formation of the final recombination products during strand exchange mediated by AtRecA2 in either chloroplasts or mitochondria. Finally, in reactions containing mitochondrial AtRecA3, addition of AtRecG alone or together with mtSSB1, mtSSB2, or OSB3 resulted in barely detectable NC products, with maximum yields of 1.0%, 0.9%, and 1.0%, respectively, accompanied by a slight decrease in JM formation relative to reactions incubated without AtRecG (**Fig. 6E, F; Table S6**). In reactions containing WHY2, NC formation increased by 1.8% relative to reactions without AtRecG (**Table S6**), reaching the highest NC yield observed with AtRecG (5.2%), similar to that obtained with AtRadA (6.4%). With OSB4, JM intermediates were barely detectable (**Fig. 6E, lanes 39–43**). Thus, results indicate that in mitochondria AtRecG may be fostering AtRecA3 mediated DNA strand exchange along with WHY2, but to a lesser extent than AtRadA. Collectively, our results indicate that meanwhile AtRadA promotes branch migration of intermediate join molecules catalyzed by plant AtRecAs helping effective DNA strand exchange in homologous recombination, AtRecG promotes accumulation of branches intermediates, which has been reported previously with bacterial RecG as possible mechanism of HR regulation (66,67).

### Combined actions of AtRadA and AtRecG helicases reveal distinct RecA dependent-HR pathways in organelles

Given that Whirlies stimulate end-product formation initiated by all three plant RecA homologs, we investigated the individual and combined effects of Whirlies, OSBs, and mtSSBs on homologous recombination in the absence and in the presence of AtRadA and AtRecG. Strand exchange reactions were incubated for 240 min, and the concentrations of all nine SSBs proteins were established with basis on their abundance measured by proteomics (68) (**Fig. 7**). Reactions incubated with AtRecA1 in the presence of WHY1 and WHY3 convert 49.8 % of the substrate to nicked circle (NC) products, the addition of OSB2 and OSB3 generated 62 % of joint molecules (JMs) that are trapped in the gel, but the combined effect of whirlies and OSBs in same reaction produce 14 % of both NC and JMs (**Fig. 7A, lane 4 to 6**). The addition of AtRadA stimulated branch migration of JMs but does not stimulate the formation of the full strand exchange product NC (**Fig. 7A, lane 7 to 9**). Surprisingly, and in contrast to the reactions described above, AtRecG stimulated NC production in reactions containing OSB2 or OSB3, while the combined action of AtRadA and AtRecG converted nearly all JM intermediates generated in the presence of OSB2 or OSB3 into NC products (**Fig. 7A, lane 10 to 15**). These results suggest that in chloroplast end-product recombination mediated by AtRecA1 proceeds through two distinct pathways: a WHY1/WHY3-dependent pathway that does not require AtRadA or AtRecG, and an OSB2/OSB3-dependent pathway that relies on AtRecG, however is less efficient in end-product formation. Although these pathways appear to operate independently, both require the activity of AtRadA and/or AtRecG for efficient completion of recombination. The greater accumulation of recombination end products observed in the presence of Whirly proteins indicates that the Whirly dependent pathway is more efficient than the OSB-dependent pathway.

**Fig. 7.**
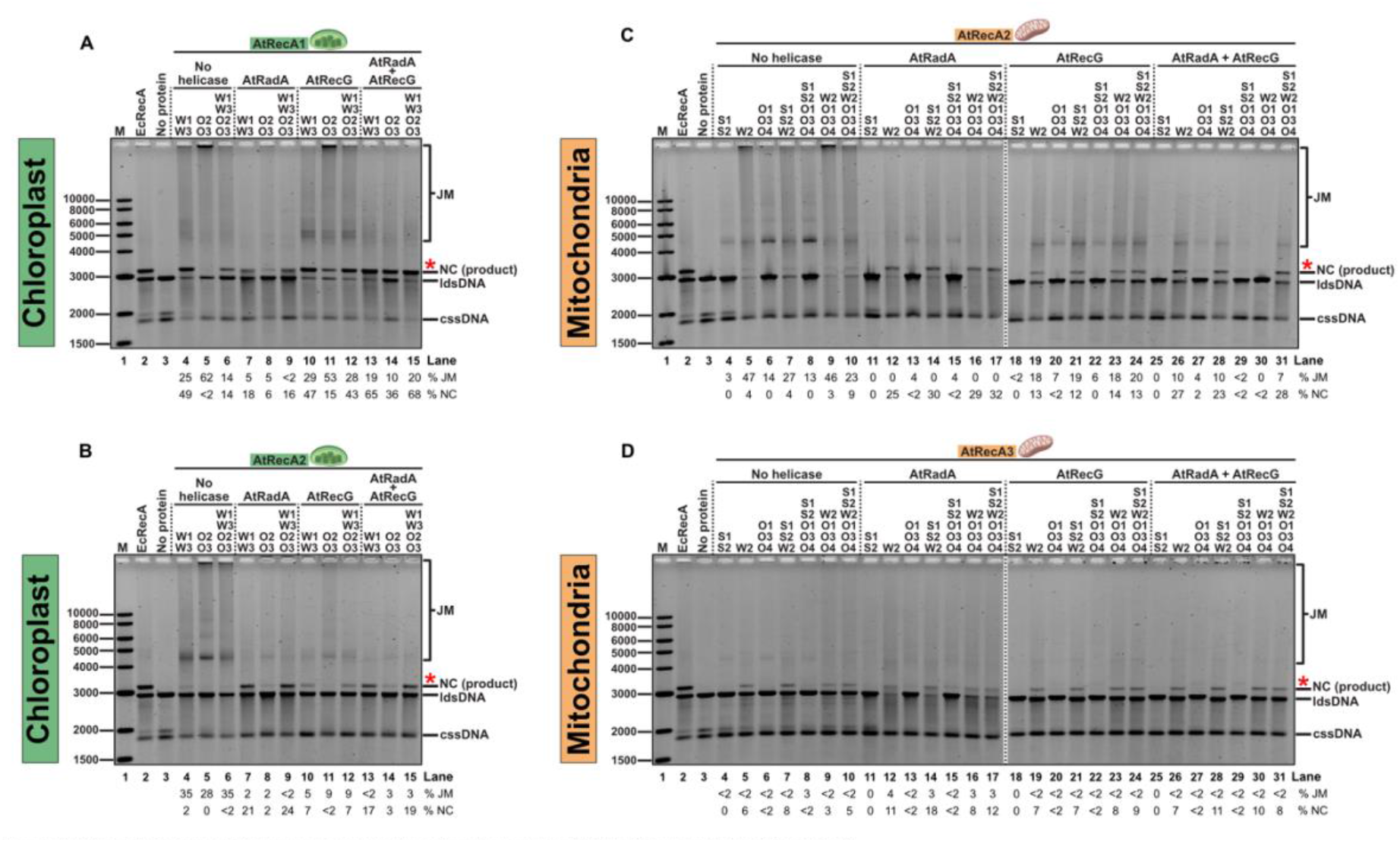
AtRadA and AtRecG synergistically promote recombination end-product formation with AtRecA1 but not with AtRecA2 or AtRecA3. (A–D) Formation of recombination end products by AtRecA1 (A), AtRecA2 (B and C), and AtRecA3 (D) in the absence of DNA helicases and in the presence of AtRadA, AtRecG, or both proteins together, along with their corresponding SSB proteins. Reactions were assembled using standard protocol, 10 μM nt cssDNA was preincubated with 3.3 μM AtRecA for 10 min, SSB proteins were added in concentration relation based on a reported mitochondrial proteomics (Fuchs et al 2020) and incubation continued for 10min. For initiating reaction 20 μM nt ldsDNA was added with 250 nM AtRadA or AtRecG helicase, and reactions were stopped after 240 min. Single stranded DNA binding protein concentrations used were: 2.5 μM Whirlies, 1.25 μM SSB1 and SSB2, 0.5 μM of OSB3 or OSB4, and 15 nM OSB1. Electrophoresis and gel staining were followed as describe in methods. (cssDNA) circular ssDNA, (ldsDNA) linear dsDNA, (NC) nicked-circle final product and (JM) joint molecules intermediate product. The amounts of NC and JM products are indicated as the percentage of total substrate added.

Regarding dually targeted AtRecA2, reactions containing WHY1 and WHY3 showed only a minimal increase in NC end-product formation (2%) and reactions containing OSB2 or OSB3 did not produce detectable NC products, although both proteins promoted JM formation (**Fig 7, lane 4 to 6**). Addition of AtRadA is required to stimulate 21-24 % of NC formation but only when both WHY1 and WHY3 are present (**Fig 7, lane 7 to 9**), and the presence of AtRecG decreased JM while minor amounts of NC are produced (7 %) when whirlies are present (**Fig 7, lane 10 to 12**). Addition of AtRadA and AtRecG (**Fig 7, lane 13 to 15**) does not induce a positive effect on NC formation in comparison to reactions in which only AtRadA is present. Again, these results suggest that, in chloroplasts, strand exchange mediated by dually targeted AtRecA2 requires both Whirlies and AtRadA as accessory proteins to generate NC products, although the reaction is less efficient than that mediated by AtRecA1.

The same analysis using AtRecA2 and mitochondrial SSB proteins, shows that in the absence of helicases, only trace amounts of NC end-product were detected in reactions containing WHY2 (4-9 %). The addition of mtSSB1 and mtSSB2 or OSBs induce the production of JM recombination intermediates, but not end-products (**Fig. 7, lanes 3 to 10**). Upon the addition of AtRadA, NC end-products are observed solely in reactions in which WHY2 is present with a 32 % maximum yield, and JMs are barely detectable (**Fig. 7, lanes 11 to 17**). An identical trend is observed in the presence of AtRecG, but the maximum yield of NC end-products was 14 %, and JMs conform 7-20 % of the products (**Fig. 7, lanes 18 to 24**). The addition of both AtRadA and AtRecG stimulates the apparition of NC end-products (**Fig. 7, lanes 25 to 31**), but JMs are produced in less than 10 %. Notably, in this assay mtSSB1 and mtSSB2 inhibited strand exchange reactions when DNA helicases are present, and even in the absence of helicases only 3% of JM is detected, which is a lower than in reactions with AtRecA2 alone (7.2% of JM) (**Fig 2E and 2F**). Thus, our data shows a similar trend to that observed in chloroplast, the minimal set for NC end-product formation in plant mitochondria mediated by AtRecA2 requires WHY2 and AtRadA helicase. Interestingly, when both AtRadA and AtRecG were present, AtRecG partially inhibited end-product formation, suggesting that the activities of these two helicases are not synergistic.

In reactions containing the mitochondria-specific AtRecA3 and WHY2, substantial levels of recombination end products were observed (**Fig. 7D, lanes 3 to 10**). The addition of AtRadA or AtRecG did not significantly enhance NC end-product formation (**Fig. 7D, lanes 11 to 31**), indicating that AtRecA3-mediated recombination is largely independent of these branch-migration helicases. Furthermore, the inclusion of organellar single-stranded DNA-binding proteins, including SSB1, SSB2, and OSB1–3, failed to further stimulate recombination beyond that achieved by AtWHY2 alone. Just as with AtReA2, SSB1 and SSB2 together inhibited strand exchange reactions when helicase is present (**Fig. 7D, lanes 11, 18 and 25**). Collectively, these results suggest that AtRecA3 primarily operates through a WHY2-dependent pathway that efficiently promotes recombination end-product formation without requiring substantial contributions from AtRadA or AtRecG. This contrasts with AtRecA1 and AtRecA2, whose activities are strongly dependent on accessory branch-migration factors and highlights a distinct mechanism of recombination processing for the mitochondria-specific AtRecA3. Thus, our analysis indicates that all organellar RecAs in combination with WHY protein induce NC end-product formation and reveal an unexpected role of whirlies in promoting homologous recombination.

### AtRecX differentially regulates AtRecA homologs, suppressing AtRecA2 while exerting minimal to no inhibitory effect on AtRecA3 and AtRecA1, respectively

Bacterial RecX is a negative regulator of RecA that limits homologous recombination by binding RecA monomers and RecA–DNA filaments, inhibiting filament extension, and actively promoting filament disassembly (69). RecX binds within the helical groove of the RecA filament, where it clamps the C-terminal domain to inhibit RecA ATPase activity and destabilize the conformational transitions required for productive filament formation and function (70–72). Plants encode a RecX ortholog with features consistent with a conserved regulatory function. In *Arabidopsis thaliana*, AtRecX is a 382-amino acid protein containing a C-terminal region that shares 45% amino acid identity with bacterial RecX proteins and an N-terminal domain of unknown function (DUF) of approximately 150 amino acids (**Fig. 8A**). Alpha-fold structural predictions suggest that the DUF N-terminal region is intrinsically disordered. Bioinformatic analyses predict that AtRecX contains a mitochondrial targeting peptide but lacks a plastid transit peptide although its subcellular localization has not yet been experimentally reported (**Fig. 8A**). The presence of AtRecX suggests that RecA activity is regulated in plant organelles. To investigate the function of RecX in HR regulation in flowering plants, we evaluated the effect of full-length AtRecX and the ΔDUF variant pn AtRecA-promoted DNA strand exchange reactions in the presence of WHY proteins and AtRadA/AtRecG helicases using AtRecX or ΔDUF variant at concentrations of 0.3 and 3.3 μM, and two different order of addition: I) AtRecX or ΔDUF variant added after incubation with WHY proteins or II) AtRecX or ΔDUF variant added 15 min after reactions were in progress (**Fig. 8B to 8E**). In reactions in which RecX was added after incubation with WHY proteins, both AtRecX and the DUF variant inhibited NC product formation at 3.3 μM, a concentration that matches the concentration of AtRecA2 and AtRecA3, whereas AtRecA1 was largely insensitive to RecX-mediated inhibition. (**Fig. 8D and 8E**). The same effect was observed when RecX was added 15 min after the reactions were in progress, AtRecA2 and AtRecA3 were inhibited by AtRecX full-length and ΔDUF variant, wherease AtRecA1 was not sensitive to RecX inhibitoty effect (**Fig. 8D and 8E**).

**Figure 8.**
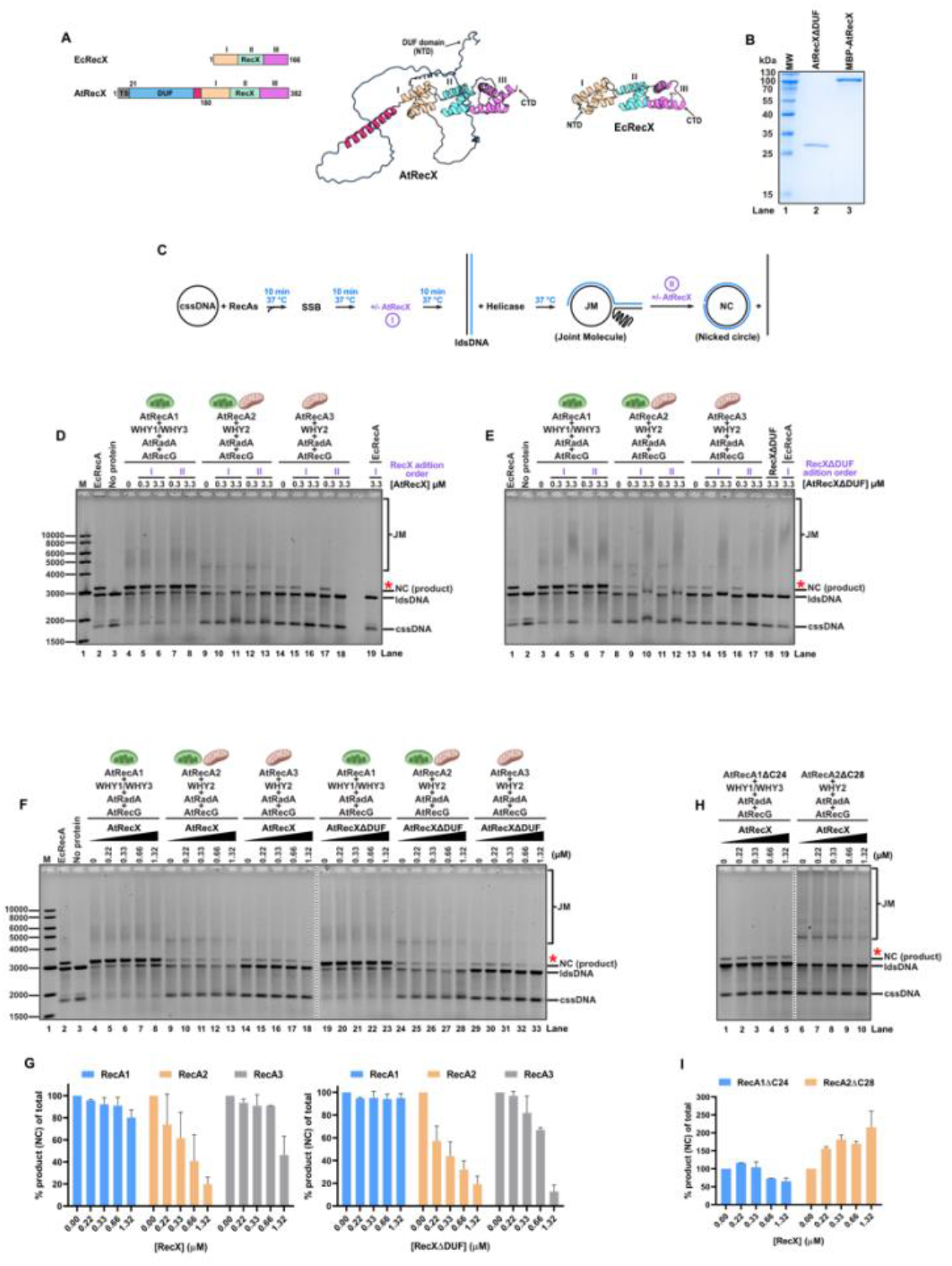
AtRecX inhibits AtRecA2-mediated strand exchange reaction. **A)** Domain organization and structural model of Arabidopsis RecX protein (AtRecX) and in comparision to RecX from *E. coli* (EcRecX). The predicted structure illustrates the position of the N-terminal DUF domain present in AtRecX relative to the conserved EcRecX core. **B)** 17 % SDS–PAGE analysis of purified AtRecX and AtRecXΔDUF proteins used in the strand-exchange assays. **C)** Schematic representation of the RecA-mediated DNA strand-exchange assay to evaluate effect of RecX. First AtRecA (3.3 μM) was preincubated with 10 μM nt cssDNA pGEM-HOM for 10 min to form the presynaptic filament, followed by addition of the corresponding SSB (2 μM) and incubation continued for 10min. AtRecX or AtRecXΔDUF was then added at the indicated concentrations. After an additional 10 min incubation, strand exchange raction was initiated by the addition of 20 μM nt ldsDNA pGEM-HOM with 250 nM of both AtRadA and AtRecG. The reactions were terminated after 120 min and run on 0.8 % agarose gel, followed by staining as described in methods, and the formation of joint-molecule (JM) intermediates and nicked-circle (NC) end products was monitored. Effect of AtRecX **(D)** or AtRecXΔDUF **(E)** order adition on strand exchange reactions mediated by AtRecA1, AtRecA2, or AtRecA3 before reactions were initiated or after 15 min reactions were in progress (I and II, respectively). **F)** Effect of AtRecX and AtRecXΔDUF on strand exchange reactions mediated by the three Arabidopsis RecA, when RecX is added before reactions were initiated (addition order I, **C**). Both AtRecX and AtRecXΔDUF inhibited the formation of recombination intermediates and final NC products to different extents among AtRecA2 and AtRecA3. **G)** Quantification of JM and NC products formed by AtRecAs in the presence of increasing concentrations of AtRecX or AtRecXΔDUF. **H)** Effect of AtRecX on a C-terminal deletion mutants of AtRecA1 and AtRecA2 on strand-exchange reactions. The C-terminal deletion of AtRecA2 substantially reduced the inhibitory effect of AtRecX on JM and NC product formation. **I)** Quantification of NC product formation in reactions containing AtRecX. Positions of cssDNA, ldsDNA, JM intermediates, and NC products are indicated.

**Figure 9.**
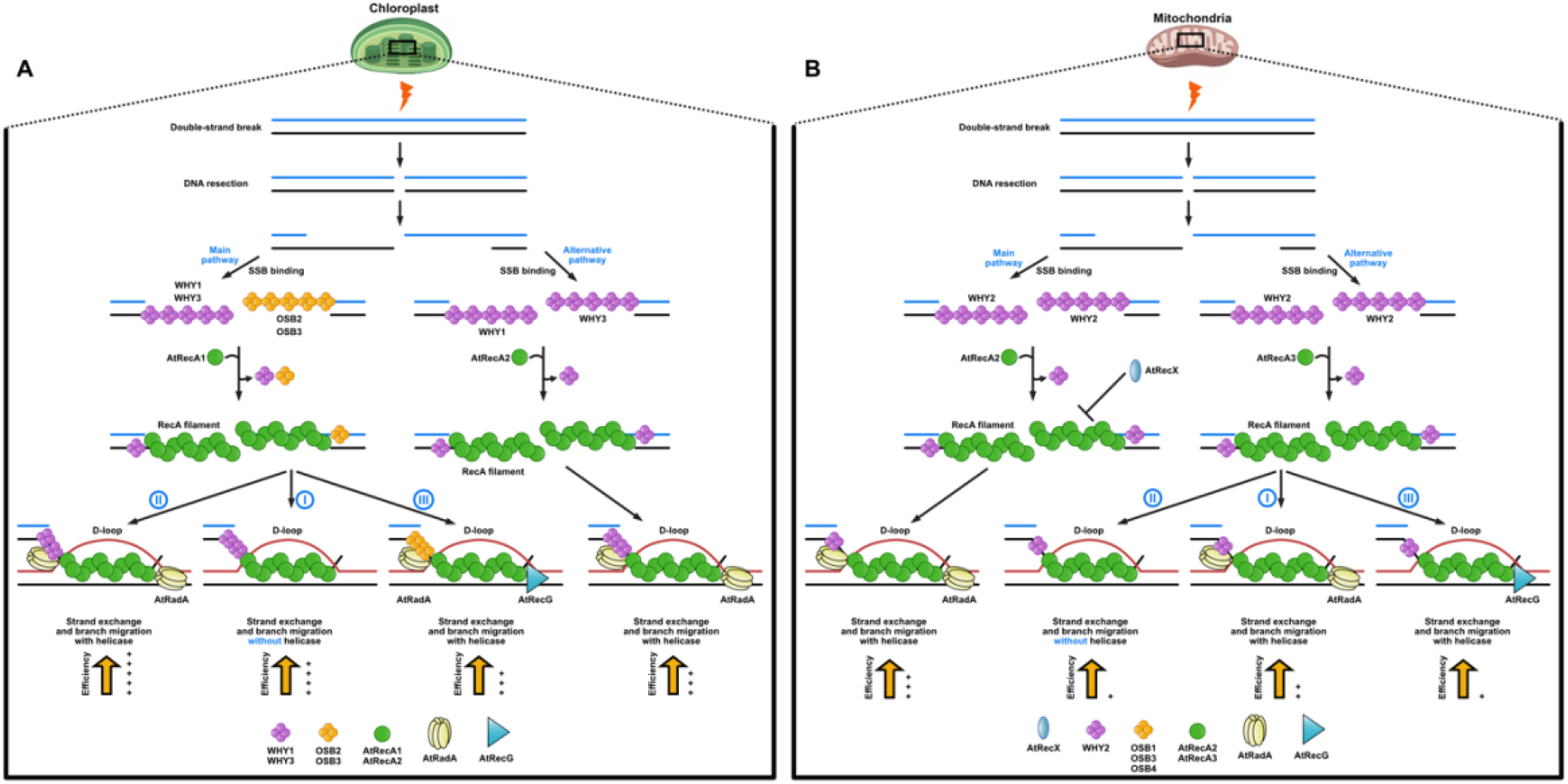
Proposed model of DNA strand exchange mediated by plant organellar recombinases. **A)** In Chloroplast, DNA strand exchnage is promoted mainly by AtRecA1. DNA-strand exchange mediated by chloroplast targeted AtRecA1 requires plant specific single stranded-DNA binding proteins WHY1/WHY3 (I), and in presence of AtRadA helicase branch migration is promoted increasing yield of NC final product. (II) Another pathway is possible with organellar single stranded-DNA binding proteins OSB2 and OSB3, and with both helicases AtRadA and AtRecG (III). An alternative pathway with dual-targeted AtRecA2 is indispensable accesory proteins WHY1/WHY3 and AtRadA helicase to promote DNA strad exchange in chloroplast. **B)** In mitochondria DNA strand exchnage is promoted mainly by AtRecA2. Dual-targeted AtRecA2 requires accesory proteins WHY2 and AtRadA helicase to promote DNA strad exchange. An alternative pathway is possible with AtRecA3 which operates through a WHY2-dependent pathway (II) that promotes recombination end-product formation, with minor required contributions from AtRadA (I) or AtRecG (III).

Next, we assessed the inhibitory effects of AtRecX and the ΔDUF variant at lower concentrations when added before the reactions were initiated (**Fig. 8F**). In reactions incubated with AtRecA1, full-length AtRecX did not inhibit nicked-circle final product at concentrations ranking from 0.22 to 0.66 µM and only 20% of inhibition was observed at 1.32 µM (RecX: RecA ratio of 1:2.5). In contrast ΔDUF variant did not significantly inhibit NC product even at 1.32 µM concentration (5 % of inhibition) (**Fig. 8F and 8G**). In reactions mediated by AtRecA2, both full-length AtRecX and the ΔDUF variant inhibited NC product formation in a dose-dependent manner, reducing the yield by up to 80% at 1.32 μM (RecX: RecA ratio of 1:2.5) (**Fig.8F and 8G**). AtRecA3-mediated reactions were more resistant to inhibition, with little effect observed at RecX concentrations up to 0.66 μM (RecX ratio of 1:5); NC product formation was reduced by up to 5% with full-length AtRecX and 33.5% with the ΔDUF variant (**Fig.8F and 8G**). This result suggests that while AtRecA1 escapes of RecX inhibition, AtRecA3 is more resistance than AtRecA2 to the inhibitory effect of AtRecX, and the RecX’s DUF domain is no relevant in this inhibition activity on AtRecAs.

Previous studies with *E. coli* RecA and the C-terminal E343K mutant showed that 3–4-fold higher concentrations of RecX were required to inhibit DNA strand exchange to a similar extent as wild-type RecA. Similarly, deletion of the last 17 amino acids reduced its sensitivity to RecX-mediated inhibition of DNA strand exchange (69). AtRecA1 harbors a C-terminal extension that contains near six continuous serines and six continuous glutamic or aspartic acids, whereas the C-terminal extension of AtRecA2 is abundant in negatively charged amino acids but the last fourteen are hydrophobic. The ability of AtRecX to inhibit after nucleoprotein filament assembly suggests that its mechanism extends beyond competition for ssDNA and likely involves direct interaction with filaments composed of AtRecA2 and AtRecA3, but not AtRecA1. To further corroborate this hypothesis, we constructed two C-terminal deletions of AtRecA1 and AtRecA2 of 24 and 28 amino acids, respectively (**Fig. S3**), as those RecAs, but not AtRecA3, harbor extended C-terminal extension (**Fig. 1A and 1B**). In reactions incubated with AtRecA1-ΔC24, only at high AtRecX concentrations of 0.66 µM and 1.32 µM up to 27.5 % and 35 % of nicked-circle product formation was reduced, respectively. In contrast, in reactions containing AtRecA2-CΔ28, AtRecX not only fails to inhibit the formation of the final nicked product but promotes the accumulation of nicked-circle products by 2-fold and JMs decreased up to 80 % (**Fig. 8H and 8I**). These results suggest that AtRecA2 C-terminal interacts with AtRecX. Together, these findings demonstrate that AtRecX differentially regulates the activities of Arabidopsis organellar RecA paralogs through a mechanism that is independent on its N-terminal DUF. The differential inhibition of the mitochondrial RecA paralogs suggests that AtRecX limits homologous recombination by restraining RecA-dependent strand exchange, specifically by AtRecA2 and suggests that AtRecX may influence the balance between homologous recombination and other DNA repair routes.

## DISCUSSION

Herein we present evidence that suggest a HR model of plant organelles, in this model AtRecA1 and AtRecA2 maintains constitutive homologous recombination, whereas AtRecA3 functions as an inducible DNA repair recombinase that operates when genome integrity is compromised. The essential nature of AtRecA1 and AtRecA2 is consistent with their predominant functions in chloroplasts and mitochondria, respectively. Loss of AtRecA1 causes embryo lethality, whereas AtRecA2 is required beyond the seedling stage, highlighting their indispensable roles in organellar genome maintenance (39,42). Chloroplast-localized AtRecA1 efficiently catalyzes homologous recombination, and accessory factors potentiate this activity, is consistent with a relatively self-sufficient recombination pathway operating in plastids. AtRecA2 relies primarily on AtRadA to promote nicked-circle formation, while AtRecG plays only a minor role in stimulating end-joining products. Thus, the absence of RadA compromises the mitochondrial function of AtRecA2, which may be partially compensated for by AtRecA3. However, the simultaneous loss of both proteins abolishes homologous recombination, ultimately leading to cell death. In contrast, AtRecA3 is dispensable under normal growth conditions but becomes essential under genotoxic stress, raising the long-standing question of why flowering plants evolved a third bacterial-type recombinase (36). This observation is consistent with the double mutant *recG recA3* exhibits pronounced developmental defects, including reduced growth, abnormal leaf morphology, variegation, and sterility, whereas *radA recA3* is lethal (26,36).

One of the most surprising findings of this study is that OSBs and Whirlies actively determine the outcome of homologous recombination rather than merely protecting ssDNA. Unlike bacterial SSB, which promotes RecA filament assembly and stimulates homologous recombination (37,73), the canonical mitochondrial SSBs (mtSSB1 and mtSSB2), do not potentiate RecA-mediated strand exchange. In contrast, whirlies and OSB1, OSB2, OSB3, and OSB4 proteins act as recombination regulators in a highly pathway-specific manner.

WHY1 and WHY3 stimulate AtRecA1-mediated strand exchange, enabling efficient formation of the final nicked circular product even in the absence of branch migration helicases. This finding provides a biochemical explanation for the relatively mild plastid phenotypes of *radA* and *recG* mutants. The mechanism by which Whirly proteins stimulate branch migration remains unclear but may involve stabilization of recombination intermediates or facilitation of DNA unwinding. Similar functional cooperation between recombinases and their cognate ssDNA-binding proteins has been described for *Borrelia burgdorferi* RecA and bacteriophage HK620 Sak4 (74,75). *why1 why3 reca1* mutants exhibit severe developmental defects and accumulate extensive chloroplast genome rearrangements, consistent with a shift towards error-prone, RecA-independent repair pathways. The notion is supported by the limited effect of *radA* defective plants on plastid HR (36). Consistent with this framework, addition of AtRadA, accelerates branch migration of recombination intermediates *in vitro*, leading to earlier formation of the final nicked circular product. It is interesting to notice that RecG and RadA mutants have limited effects on the stability of the plastidial genome (26,36) a situation that correlates with the formation of final nicked-circle products in the sole presence of AtRecA1 and chloroplast whirlies.

Our results indicate that the dual-targeted recombinase AtRecA2 displays an even greater degree of specialization, as AtRecA2 supports homologous recombination in both chloroplasts and mitochondria through distinct sets of accessory ssDNA-binding proteins. In chloroplasts, WHY1 and WHY3, promote AtRecA2-mediated recombination, whereas in mitochondria this role is fulfilled primarily by WHY2, but AtRecA2 strand exchange activity depends on both whirlies and AtRadA to promote nicked-circle formation into chloroplast and mitochondria. These findings reveal that the same recombinase can operate within different molecular contexts depending on the organelle in which it functions, illustrating how plant-specific ssDNA-binding proteins have diversified to tailor homologous recombination to the distinct physiological requirements of chloroplasts and mitochondria. Thus, our biochemical reconstitution establishes Whirly proteins as *bona fide* accessory recombination factors that actively promote high-fidelity homologous recombination rather than serving solely as ssDNA-binding proteins.

The mitochondria-specific AtRecA3 defines a third homologous recombination pathway. In our reconstituted system, efficient completion of strand exchange occurred only in the presence of WHY2, identifying this protein as the principal accessory factor for AtRecA3. Unlike AtRecA2, AtRecA3 exhibited little dependence on AtRadA or AtRecG, suggesting that it operates through a mechanistically distinct pathway specialized for mitochondrial DNA repair. However, the *radA recA3* is lethal, indicating that RECA3 is an alternative DNA repair route in plant mitochondria (36). Seminal work from Gualberto’s group demonstrated that, in mitochondria, RadA functions as the primary branch-migration pathway (36). AtRadA strongly stimulates AtRecA1 NC-formation, particularly in the presence of WHY1 and WHY3. Notably, AtRecA2-mediated end-product formation depends on the presence of AtRadA, whereas chloroplast-localized AtRecA1 only generate nicked products in its absence. Together, these results indicate that dual-targeted AtRecA2 relies on both RadA and Whirly proteins, whereas RadA function is not essential for chloroplast AtRecA1. In sum, our work shows that Whirlies stimulate NC formation, whereas OSB1 and OSB3 causes NC inhibition or JM accumulation, respectively, this SSBs proteins differentially regulate HR regulation. Similarly, mtSSB1 and mtSSB2 inhibited strand exchange (**Fig. 7C and 7D**) in strand exchange reaction coupled with DNA helicases.

Although AtRecG exhibits branch migration activity and promotes nicked-circular (NC) product formation from isolated branched intermediates, it inhibits NC formation and promotes junction molecule (JM) accumulation in reactions coupled with RecA filament formation. A similar effect has been reported for bacterial RecG and is attributed to the opposite directionality of RecG-mediated branch migration relative to RecA-mediated strand exchange, resulting in reverse branch migration (66).

An additional level of regulation is provided by AtRecX, whereas bacterial RecX acts as a universal antagonist of RecA filament assembly, Arabidopsis RecX exhibits pronounced paralog specificity. AtRecX primarily suppresses strand exchange catalyzed by AtRecA2. This paralog-specific regulation likely provides a mechanism for fine-tuning homologous recombination. Consistent with this hypothesis, altered RECX expression in *Physcomitrella* results in mitochondrial elevated ectopic recombination between short, dispersed repeats, whereas chloroplast recombination remains largely unaffected despite the reported plastid localization of the protein, suggesting that RecX restrains RecA activity to promote accurate homologous recombination and preserve mitochondrial genome integrity (76).

Finally, a direct compassion between the final yield of nicked-circular (NC) products in DNA strand exchange reactions mediated by plant organellar recombinases, AtRecA1 exhibits yields comparable to those reported for bacterial recombinases In contrast, AtRecA2 and AtRecA3 produce NC yields comparable to those reported for the yeast recombinase Rad51, which requires three accessory proteins to promote NC formationA (29,77–79).

Collectively, our findings demonstrate that homologous recombination in plant organelles is not a single conserved bacterial pathway, but a collection of specialized recombination systems assembled from proteins of diverse evolutionary origins. Bacterial-derived proteins, including RecA, RadA, RecG, and RecX, provide the catalytic core, whereas plant-specific Whirly and OSB proteins determine pathway choice and recombination efficiency in a compartment-specific manner. Together with surveillance factors such as MSH1 that suppress ectopic recombination, these proteins form a multilayered network that enables plant mitochondria and chloroplasts to maintain genome stability while preserving the remarkable genomic plasticity that characterizes plant organellar DNA evolution. Our findings underscore that plant organellar homologous recombination is a mosaic process integrating bacterial, phage-derived, and plant-specific activities to preserve genome stability while permitting controlled genomic plasticity.

## AUTHOR CONTRIBUTIONS

JM-G: Investigation, Formal analysis, Methodology, Validation, Writing—original draft. APC: Formal analysis, visualization, Methodology. AAG: Investigation, Formal analysis, Methodology. CDR: Investigation, Formal analysis, Methodology. CDQ: Formal analysis, Methodology. LGB: Conceptualization, Founding, Writing—review & editing

## SUPPLEMENTARY DATA

Supplementary Data are available online.

## DATA AVAILABILITY STATEMENT

All data are incorporated into the article and its online supplementary material

