## Supplemental information for "Plant Organellar Homologous Recombination Depends on Accessory Factors for Recombinase Activity"

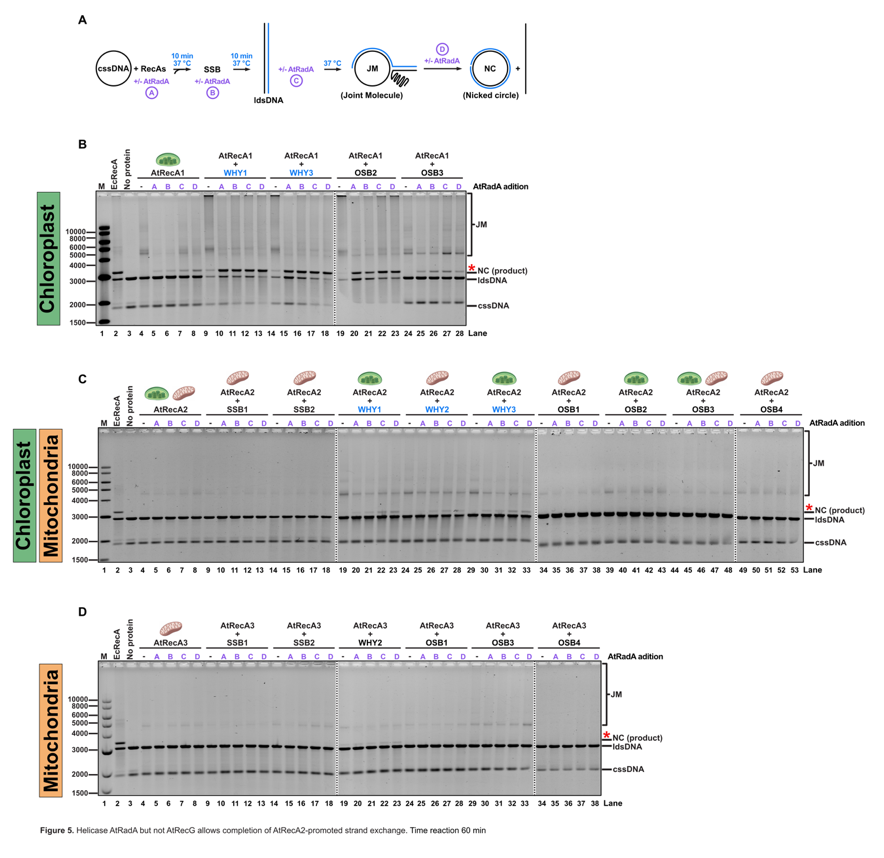

**Fig. S1. AtRadA addition order in strand exchange reactions has a slightly effect on NC product yield only with AtRecA2 and whirlies.** Four protein-addition schemes were tested to determine whether the order of addition of helicase influences the formation of nicked circular products at 60 minutes time reaction. In all cases, standard conditions described in methods were followed with circular ssDNA (cssDNA) preincubation with AtRecA, and the following addition schemes were tested: **(A)** helicase was added simultaneously with AtRecA **(B)** helicase was added simultaneously with SSB, **(C)** helicase was added simultaneously with ldsDNA, and **(D)** helicase was added 15 min after reaction was in progress.

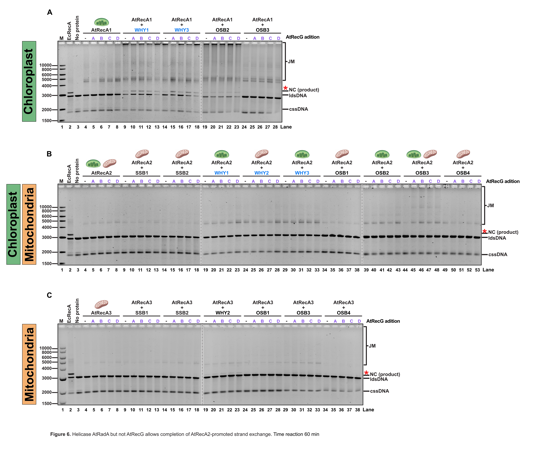

**Fig. S2. Order of AtRecG addition** **in strand exchange reactions has minimal effects on final nicked circular product formation.** Four protein-addition schemes were tested to determine whether the order of addition of helicase influences the formation of nicked circular products at 60 minutes time reaction. In all cases, standard conditions described in methods were followed with circular ssDNA (cssDNA) preincubation with AtRecA, and the following addition schemes were tested: **(A)** helicase was added simultaneously with AtRecA **(B)** helicase was added simultaneously with SSB, **(C)** helicase was added simultaneously with ldsDNA, and **(D)** helicase was added 15 min after reaction was in progress. The order of protein addition had minimal effect on the formation of the final nicked circular product in reactions containing AtRecG.

| **TableS1. Plasmid constructions.** | | | | | |
| --- | --- | --- | --- | --- | --- |
| **Gene** | **Accession**  **number** | **Expression vector** | **Restriction site** | **Primer number** | **Primers used for cloning or mutagenesis**  **(5’ → 3’)** |
| SSB1 | At4g11060 | pET19b | *Nde*I/  *Bam*HI | (1) Fw  (2) Rv | aagatcacatatgaagtcgtggttctctactggtc  ggttcttggatccctaaatcaatccttcttttagc |
| SSB2 | At3g18580 | pCri1b | *Nde*I/  *Bam*HI | (3) Fw  (4) Rv | aagatcacatatgcaagcttccatgtctttctgtac  ggttcttggatcctcagtagtagccaacgcctc |
| WHY1 | AT1G14410 | pET19b | *Nde*I/  *Bam*HI | (5) Fw  (6) Rv | aaggaaccatatgccttcacctgctgaaggattgcct  tcggtccggatcctcatctattccattcatagtctcc |
| WHY2 | At1g71260 | pET28b | *Nde*I/  *Sac*I | (7) Fw  (8) Rv | aagatcacatatgaaagatgccgcaaagcctagtg  ggttcttgagctctcatttatcccactccagctc |
| WHY3 | AT2G02740 | pET19b | *Nde*I/  *Bam*HI | (9) Fw  (10) Rv | aaggaaccatatgaacgctgaagtatcttctcctaga  tcggtccggatcctcatctactccattcgtagtctc |
| OSB1 | At1g47720 | pCri1b | *Nde*I/  *Bam*HI | (11) Fw  (12) Rv | aagatcacatatgcatgcgcgtttgttcaagaaacc  ggttcttggatccctaaatatagtcagaaagacg |
| OSB2 | At4g20010 | pET19b | *Nde*I/  *Bam*HI | (13) Fw  (14) Rv | aagatcacatatggtgaaaccgctggagattgcgtc  ggttcttggatccctaaaagacaatactttcttgtttc |
| OSB3 | At5g44785 | pET19b | *Nde*I/  *Bam*HI | (15) Fw  (16) Rv | aagatcacatatgccaccgctaaatgtagcgacggag  ggttcttggatccttaggccatgacaggtctttcttg |
| OSB4 | At1g31010 | pET28b SUMO | *Nde*I/  *Bam*HI | (17) Fw  (18) Rv | cgccgccatatgctttcaacttcttctacggaaag  cgcctcggatccttagaacatctcttgtacaccgt |
| AtRecA1 | AT1G79050 | pCri1b | *Nde*I/  *Bam*HI |  | Synthetic gene cloning from pTwist plasmid with restriction enzymes |
| AtRecA1ΔC24 |  |  |  | (19) Fw  (20) Rv | TTCAACGCCGtaaATGTCGTCGA  TCGACGACATttaCGGCGTTGAA |
| AtRecA2 | At2g19490 | pCri1b | *Nde*I/  *Bam*HI |  | Synthetic gene cloning from pTwist plasmid with restriction enzymes |
| AtRecA2ΔC28 |  |  |  | (21) Fw  (22) Rv | TAAGGATCCGAATTCGAG  CAGGCTGTCCTCTTC |
| AtRecA3 | At3g10140 | pCri1b | *Nde*I/  *Bam*HI |  | Synthetic gene cloning from pTwist plasmid with restriction enzymes |
| AtRecA3  K463P |  |  |  | (23) Fw  (24) Rv | GGTGTACGGAccgGAAGCCTCGG  TCTACCATGCGTCCTTTTG |
| AtRecA3  P615K |  |  |  | (25) Fw  (26) Rv | AAACGCTTTGaaaTTCCACGCAGC  CCGCCACATGTGACC |
| AtRecA3  K463P, P615K |  |  |  | (N) Fw  (N) Rv | Doble mutant was made from single mutant |
| AtRecX | At3g13226.1 | pCri1b | *Nde*I/  *Bam*HI |  | Synthetic gene cloning from pTwist plasmid with restriction enzymes |
| AtRecXΔDUF |  |  | *Nde*I/  *Bam*HI | (27) Fw  (28) Rv | ggaggaacatatgcacaagcaagcctatgataagatc  gaatgccggatcctcaggattcgtgggt |
| AtRadA  (ΔN88) | At5g50340 | pET19b | *Nde*I/  *Bam*HI | (29) Fw  (30) Rv | ggaggagcatatggtcgcctccgttttcgacctt  ttaataaggatccttatcccttaaacacagcattgat |
| AtRecG  (ΔN56) | At2g01440.1 | pCri1b | *Nde*I/*Hind*III | (31) Fw  (32) Rv | ccaccaacatatggacaatcttcttgagcaagttgag  ttaataaaagcttttagtcgcccagaagacaaagtgg |

| **Table S2.** | |
| --- | --- |
| **Protein** | **MW**  **(kDa)** |
| SSB1 | 19.63 |
| SSB2 | 21.91 |
| WHY1 | 21.81 |
| WHY2 | 21.95 |
| WHY3 | 21.74 |
| OSB1 | 26.62 |
| OSB2 | 34.86 |
| OSB3 | 44.38 |
| OSB4 | 33.87 |
| MBP-AtRecA1 | 87.11 |
| MBP-AtRecA1ΔC24 | 84.42 |
| MBP-AtRecA2 | 86.41 |
| MBP-AtRecA2ΔC28 | 84.00 |
| MBP-AtRecA3 | 83.37 |
| MBP-AtRecA3  K463P | 83.34 |
| MBP-AtRecA3 P615K | 83.40 |
| MBP-AtRecA3  K463P, P615K | 83.50 |
| MBP-AtRecX | 87.89 |
| AtRecXΔDUF | 23.21 |
| AtRadA  (ΔN88) | 61.58 |
| MBP-AtRecG  (ΔN56) | 148.11 |

| **Table S3. Oligonucleotides used for assays** | | | |
| --- | --- | --- | --- |
| **Biochemical assay** | **(Number)**  **name** | **Length**  **(nt)** | **Oligonucleotide sequence**  **(5’ → 3’)** |
| ATPase | (1)  Cox2-RP | 57 | TGCCCTTCTTCTAGCAGCAGTCTTCTTATATACGAAACCAACATCCTTATAATACTA |
|  | (2)  Cox2-EcoRI-3 | 27 | TAGTATTATAAGGATGTTGGTTTCGTA |
| D-loop | (3)  M13 | 90 | TTTTGCAAAAGAAGTTTTGCCAGAGGGGGTAATAGTAAAATGTTTAGACTGGATAGCGTCCAATACTGCGGAATGTCATAAATATTCAT |

| **Table S4** | | | | | | | | | | |
| --- | --- | --- | --- | --- | --- | --- | --- | --- | --- | --- |
| AtRecA1 (240 min) | | | | | | | | | | |
|  | Without helicase | | AtRadA | | % change in yield | | AtRecG | | % change in yield | |
|  | % NC | % JM | % NC | % JM | NC | JM | % NC | % JM | NC | JM |
| No SSB | 2.5 | 53.4 | 13.7 | 2.7 | +11.2 | -50.7 | 3.9 | 55.9 | +1.4 | +2.5 |
| **WHY1** | 41.6 | 53.1 | 57 | 29 | **+15.4** | **-24.1** | 23.4 | 72.2 | **-18.2** | **+19.1** |
| **WHY3** | 51.3 | 42.9 | 67.5 | 16.5 | **+16.2** | **-26.4** | 33.2 | 61.1 | **-18.1** | **+18.2** |
| **OSB2** | 15 | 83 | 42 | 33 | **+27** | **-50** | 21.1 | 75.3 | +6.1 | -7.7 |
| **OSB3** | 0.8 | 26 | 14.7 | 44.5 | +13.9 | **+18.5** | 4.8 | 58.3 | +4 | **+32.3** |

| **Table S5.** | | | | | | | | | | |
| --- | --- | --- | --- | --- | --- | --- | --- | --- | --- | --- |
| AtRecA2 (240 min) | | | | | | | | | | |
|  | Without helicase | | AtRadA | | % change in yield | | AtRecG | | % change in yield | |
|  | % NC | % JM | % NC | % JM | % NC | % JM | % NC | % JM | % NC | % JM |
| No SSB | 0.0 | 7.6 | 1.1 | 1.2 | +1.1 | -6.4 | 0.0 | 5.5 | 0.0 | -2.1 |
| SSB1 | 0.0 | 5.6 | 0.6 | 1.0 | +0.6 | -4.6 | 0.0 | 6.0 | 0.0 | +0.4 |
| SSB2 | 0.0 | 5.5 | 0.7 | 1.1 | +0.7 | -4.4 | 0.0 | 11.7 | 0.0 | +6.2 |
| **WHY1** | 1.9 | 48.3 | 12.0 | 6.0 | **+10.1** | **-42.3** | 0.8 | 41.4 | -1.1 | -6.9 |
| **WHY2** | 0.5 | 60.3 | 15.5 | 25.1 | **+15.0** | **-35.2** | 1.7 | 63.2 | +1.2 | +2.9 |
| **WHY3** | 1.6 | 41.2 | 13.8 | 4.5 | **+12.2** | **-36.7** | 1.0 | 31.6 | -0.6 | -9.6 |
| **OSB1** | 0.0 | 0.7 | 0.6 | 12.3 | +0.6 | **+11.6** | 0.0 | 6.0 | 0.0 | +5.3 |
| OSB2 | 0.0 | 11.8 | 3.7 | 6.4 | +3.7 | -5.4 | 0.0 | 16.6 | 0.0 | +4.8 |
| **OSB3** | 0.0 | 4.3 | 4.9 | 30.1 | +4.9 | **+25.8** | 0.0 | 44.9 | **0.0** | **+40.6** |
| OSB4 | 0.0 | 3.9 | 1.0 | 3.2 | +1.0 | -0.7 | 0.0 | 7.4 | 0.0 | +3.5 |

| **Table S6.** | | | | | | | | | | |
| --- | --- | --- | --- | --- | --- | --- | --- | --- | --- | --- |
| AtRecA3 (240 min) | | | | | | | | | | |
|  | Without helicase | | AtRadA | | % change in yield | | AtRecG | | % change in yield | |
|  | % NC | % JM | % NC | % JM | % NC | % JM | % NC | % JM | % NC | % JM |
| No SSB | 0.0 | 1.5 | 0.0 | 0.2 | 0.0 | -1.3 | 1.0 | 6.0 | 1.0 | +4.5 |
| SSB1 | 0.0 | 0.7 | 0.0 | 0.0 | 0.0 | -0.7 | 0.9 | 0.5 | 0.9 | -0.2 |
| SSB2 | 0.0 | 0.9 | 0.38 | 0.0 | **0.38** | -0.9 | 1.0 | 0.5 | 1.0 | -0.4 |
| **WHY2** | **3.4** | 2.8 | **6.4** | 0.4 | **3.0** | **-2.4** | **5.2** | **4.1** | **1.8** | **+1.3** |
| OSB1 | 0.0 | 2.0 | 0.0 | 0.7 | 0.0 | -1.3 | 0.0 | 1.3 | 0.0 | -0.7 |
| **OSB3** | 1.4 | 7.0 | 1.5 | 21.7 | **0.1** | **+14.7** | 2.9 | 5.0 | **1.5** | **-2.0** |
| OSB4 | 0.0 | 1.5 | 0.0 | 0.3 | 0.0 | -1.2 | 0.0 | 0.8 | 0.0 | -0.7 |
